# TDP43 autoregulation gives rise to shortened isoforms that are tightly controlled by both transcriptional and post-translational mechanisms

**DOI:** 10.1101/2024.07.02.601776

**Authors:** Megan M. Dykstra, Kaitlin Weskamp, Nicolás B. Gómez, Jacob Waksmacki, Elizabeth Tank, M. Rebecca Glineburg, Allison Snyder, Emile Pinarbasi, Michael Bekier, Xingli Li, Jen Bai, Shameena Shahzad, Juno Nedumaran, Clare Wieland, Corey Stewart, Sydney Willey, Nikolas Grotewold, Jonathon McBride, John J. Moran, Aditya V. Suryakumar, Michael Lucas, Peter Tessier, Michael Ward, Peter Todd, Sami J. Barmada

## Abstract

The nuclear RNA-binding protein TDP43 is integrally involved in the pathogenesis of amyotrophic lateral sclerosis (ALS) and frontotemporal lobar degeneration (FTLD). Previous studies uncovered N-terminal TDP43 isoforms that are predominantly cytosolic in localization, highly prone to aggregation, and enriched in susceptible spinal motor neurons. In healthy cells, however, these shortened (s)TDP43 isoforms are difficult to detect in comparison to full-length (fl)TDP43, raising questions regarding their origin and selective regulation. Here, we show that sTDP43 is created as a byproduct of TDP43 autoregulation and cleared by nonsense mediated RNA decay (NMD). The sTDP43-encoding transcripts that escape NMD can lead to toxicity but are rapidly degraded post-translationally. Circumventing these regulatory mechanisms by overexpressing sTDP43 results in neurodegeneration *in vitro* and *in vivo* via N-terminal oligomerization and impairment of flTDP43 splicing activity, in addition to RNA binding-dependent gain-of-function toxicity. Collectively, these studies highlight endogenous mechanisms that tightly regulate sTDP43 expression and provide insight into the consequences of aberrant sTDP43 accumulation in disease.

## Introduction

Amyotrophic lateral sclerosis (ALS) and frontotemporal lobar degeneration (FTLD) are two fatal neurodegenerative disorders lacking comprehensive treatment options. Distinct populations of neurons are vulnerable in each disease, correlating with their unique clinical presentations. Even so, the majority of those with ALS and FTLD share a signature neuropathological feature characterized by the nuclear exclusion and cytoplasmic accumulation of TDP43, an essential and typically nuclear RNA-binding protein^1–3^. Despite its prevalence, the mechanisms underlying TDP43 mislocalization, and the corresponding impact of TDP43 cytoplasmic deposition and/or loss of nuclear TDP43 in disease remain unclear.

TDP43 is a ubiquitously expressed RNA-binding protein involved in several critical steps of RNA processing. Through its two RNA recognition motifs, TDP43 binds intronic UG-rich sequences present in approximately one-third of all transcribed RNAs^4–5^, where it predominantly acts as a splicing repressor, blocking the incorporation of unannotated and non-conserved ‘cryptic’ exons into mature mRNA^6–7^. Cryptic exons have been detected in postmortem ALS and FTLD tissue as well as multiple disease models^8–13^, and thus provide a reliable readout of TDP43 dysfunction in disease. Given the extensive nature of TDP43 RNA substrates, mislocalization of this protein in disease may drive the misprocessing of thousands of RNAs, including several that are essential for neuronal function such as those encoding *STMN2*, *UNC13A,* and *KALRN*^8–9,11–12,14^, ultimately resulting in neurodegeneration.

In prior studies, we and others uncovered truncated TDP43 splice isoforms that lack the C-terminal low complexity domain^15–20^. These shortened (s)TDP43 variants are enriched in motor neurons^19^, the cell type most vulnerable in ALS, and contain a novel 18-amino acid sequence encoding a functional nuclear export sequence (NES). As a result, the distribution of sTDP43 is primarily cytoplasmic, in direct contrast to native full-length (fl)TDP43^19^. Overexpressed sTDP43 sequesters flTDP43 within cytosolic inclusions reminiscent of neuropathological changes in ALS and the subset of FTLD with TDP43 pathology (FTLD-TDP)^19^, raising the possibility that sTDP43 accumulation may underlie disease-associated loss of TDP43 splicing activity and mislocalization.

Despite sTDP43 being evolutionarily conserved^16–20^, the factors that regulate sTDP43 both physiologically and pathophysiologically remain unknown. Here, we identify nonsense-mediated mRNA decay (NMD) as the primary pathway minimizing sTDP43 production in immortalized cells and mature neurons. We also describe post-translational mechanisms that complement NMD in ensuring baseline, low levels of sTDP43. In the absence of these regulatory pathways, sTDP43 accumulation leads to neurotoxicity in part by affecting the localization and splicing activity of flTDP43. Together, these investigations may prove essential for elucidating the conserved function of sTDP43, as well as the consequences of its accumulation.

## Results

### sTDP43-encoding transcripts are NMD substrates generated by TDP43 autoregulation

Our work and that of others emphasized the critical importance of maintaining steady-state TDP43 levels in neurons; even small increases or decreases in TDP43 abundance can lead to cell death^21–37^. As such, TDP43 expression is strictly regulated through a negative feedback loop termed autoregulation^38–39^. At high levels, TDP43 binds to a UG-rich sequence located in the 3’ untranslated region (3’UTR) of its cognate mRNA^4–5^. Recognition of this sequence (called the TDP43 binding region, or TBR), prompts 3’UTR splicing and the use of alternative polyadenylation signals (**Figure 1a**). The alternatively spliced variants are retained in the nucleus and degraded by the RNA exosome, but a portion likely escapes to the cytosol, where they are degraded by NMD^38–39^.

**Figure 1:**
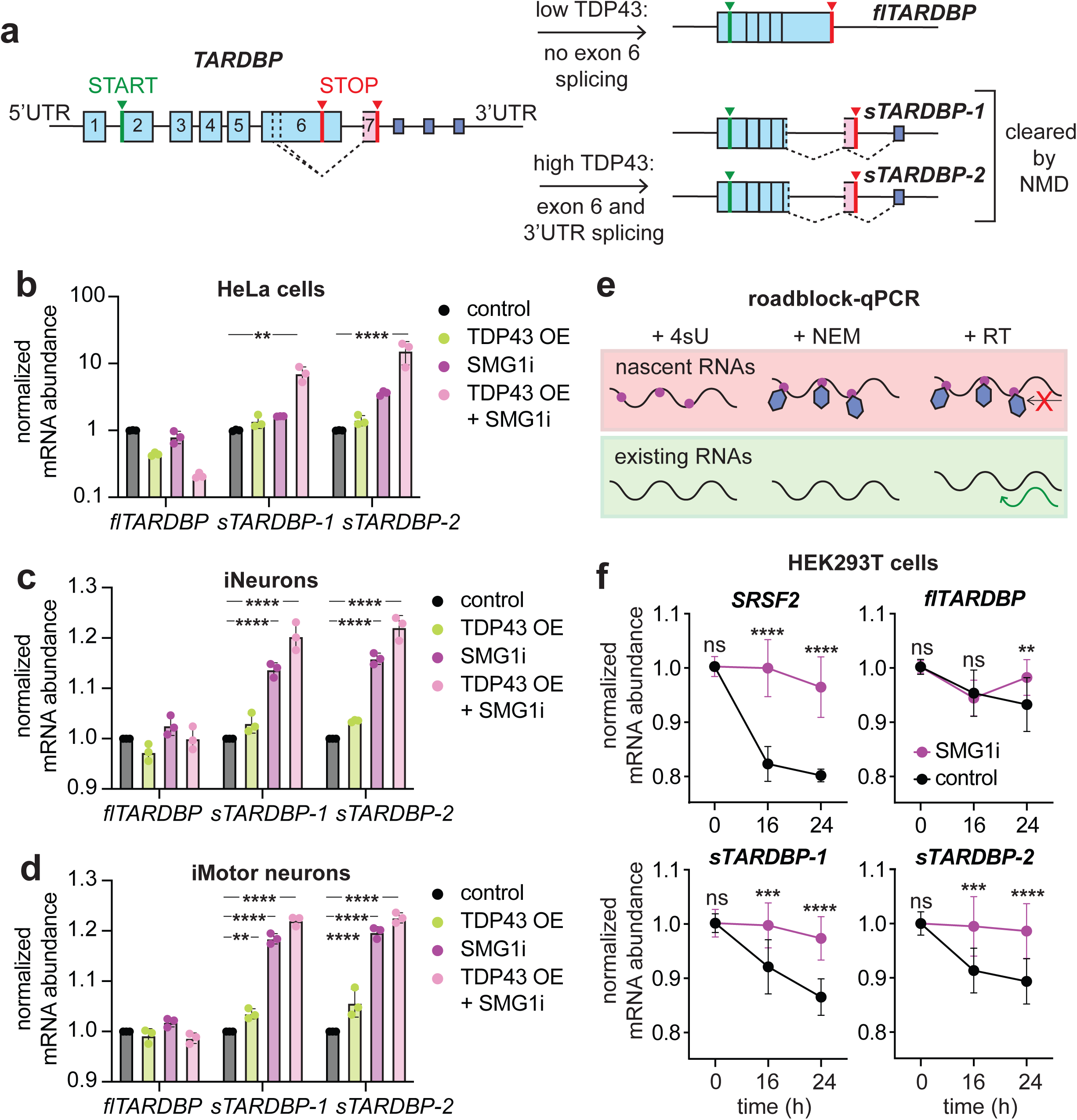
sTDP43-encoding transcripts are NMD substrates generated by TDP43 autoregulation. (**a**) Diagram depicting TDP43 autoregulation. TDP43 binds its own 3’UTR, triggering alternative splicing and the generation of sTDP43-encoding transcripts predicted to be nonsense mediated mRNA decay (NMD) substrates due to splicing downstream of termination codon. (**b**) qRT-PCR showing sTDP43 transcript abundance increases after overexpressing TDP43 (TDP43 OE) and blocking NMD using SMG1i in Flp-In GFP-TDP43 HeLa cells. SMG1i and TDP43 OE also increase sTDP43 transcript levels in human iNeurons (**c**) and iMotor neurons (**d**). (**e**) Schematic of roadblock-qPCR. (**h**) Stability of *SRSF2*, an NMD substrate, as well as *flTARDBP*, *sTARDBP-1,* and *sTARDBP-2* in HEK293T cells treated with vehicle or SMG1i. (**b**-**d**, **f**) Data collated from 3 experiments. **p<0.01, ***p<0.001, ****p<0.0001 by 2-way ANOVA with Tukey’s multiple comparisons test.

Several lines of evidence, including data from high-throughput sequencing studies and northern blotting experiments, suggest that truncated transcripts generated by TDP43 autoregulation correspond to sTDP43 variants detected in human neurons and postmortem tissue^15–16,18–20,40^. To test this, we first induced TDP43 autoregulation and measured sTDP43 RNA by quantitative (q)RT-PCR. Flp-In GFP-TDP43 HeLa cells^41^ contain a single copy of exogenous, GFP-tagged TDP43, which, upon the addition of doxycycline (DOX), is expressed at levels sufficient to induce autoregulation^4^. After 48h of TDP43 overexpression (OE), we noted a modest reduction of the full-length (fl)*TARDBP* transcript in DOX-treated cells, indicative of autoregulation (**Supplemental Figure 1a**). Under these same conditions, we observed a subtle increase in the abundance of *sTARDBP* isoforms 1 and 2 compared to vehicle-treated controls (**Supplemental Figure 1a**). These variants utilize the same splice acceptor site, but splice donor sites that differ only by nine nucleotides. To determine if sTDP43-encoding transcripts are subject to NMD, we treated Flp-In GFP-TDP43 HeLa cells with cycloheximide (CHX), a translational repressor that blocks translation-dependent processes such as NMD^42^. Confirming the ability of CHX to inhibit NMD, we noted a significant upregulation of *SRSF2*, a canonical NMD substrate^43^, in CHX-treated HeLa cells (**Supplemental Figure 1a**). CHX also increased *sTARDBP-1* and *-2*, similar to what was observed for TDP43 OE after the addition of DOX. Simultaneous treatment with DOX and CHX increased the abundance of both *sTARDBP-1* and *-2* (**Supplemental Figure 1a**), consistent with the generation of *sTARDBP* RNA by TDP43 autoregulation and their degradation through NMD.

CHX is a relatively crude NMD inhibitor with substantial cytotoxicity and off-target effects^44^. Therefore, to confirm the sensitivity of *sTARDBP* transcripts to NMD, we repeated these experiments using 11j (SMG1i), a small molecule that selectively blocks the phosphorylation of UPF1 by SMG1^45^, an essential step in the NMD pathway (**Supplemental Figure 1b**). In keeping with our prior results, the combined treatment of DOX (TDP43 OE) and SMG1i increased the abundance of *sTARDBP*-1 and -2 ten-fold (**Figure 1b**), providing further evidence for *sTARDBP* targeting by NMD.

We also investigated *sTARDBP* processing and stability in human neurons. For these experiments, we differentiated human induced pluripotent stem cells (iPSCs) from healthy donors into glutamatergic, excitatory forebrain-like iNeurons through the directed expression of the master transcription factors Ngn1 and Ngn2^19,46–49^. In accordance with the data from HeLa cells, SMG1i treatment resulted in a significant increase in *sTARDBP-1* and *-2* in iNeurons (**Figure 1c**). Similar results were obtained in iMotor neurons differentiated from iPSCs by regulated expression of Ngn2, Islet1 and Lhx3^48,50–52^ (**Figure 1d**). These data strongly indicate that *sTARDBP* transcripts are subject to NMD in multiple cell types, ranging from transformed cells to human neurons.

qRT-PCR is a sensitive method for comparing the abundance of individual RNA transcripts but provides little information on RNA clearance rates *per se*. To determine if NMD contributes directly to the turnover of *sTARDBP* mRNA, we employed roadblock-qPCR^53^, a creative technique for measuring the decay of endogenous transcripts *in cellulo* (**Figure 1e**). HEK293T cells were treated with 4-thiouridine (4sU), a uracil analog that is freely incorporated into newly synthesized transcripts. 4sU-labeled RNA was then modified by N-methylmaleimide (NEM), a bulky moiety that acts as a ‘roadblock’ to reverse transcriptase, effectively eliminating all nascent RNAs from being copied into cDNA. Roadblock-qPCR therefore enables determination of transcript half-life by following the abundance of pre-existing RNA transcripts at specific times following 4sU incorporation. For controls, we tracked the turnover of *SRSF2*, an NMD target^54^, and *flTARDBP*, which is insensitive to NMD^38^. As expected, *SRSF2* was highly unstable, but significantly stabilized by treatment with SMG1i; on the other hand, *flTARDBP* transcripts were remarkably stable and unaffected by SMG1i. Both *sTARDBP-1* and *-2* were unstable in comparison to *flTARDBP,* yet effectively stabilized by SMG1i (**Figure 1f**). Together with the data demonstrating an increase in *sTARDBP* isoforms upon TDP43 overexpression, these experiments support that sTDP43-encoding transcripts are produced via TDP43 autoregulation and suppressed via NMD.

### NMD and TDP43 autoregulation dictate sTDP43 protein levels

To determine if the observed changes in *sTARDBP* transcript abundance are mirrored by changes at the protein level, we transfected mouse neuroblastoma (N2a) cells with EGFP-TDP43 (TDP43 OE), and/or treated them with SMG1i for 48h, followed by immunoblotting (**Figure 2a**). Using an antibody that recognizes the N-terminus of TDP43—and therefore detects exogenous EGFP-TDP43, endogenous flTDP43 as well as endogenous sTDP43—we again detected increases in sTDP43 immunoreactivity following TDP43 OE and NMD inhibition (**Figure 2a**, **b**). In comparison, we detected a subtle but insignificant drop in endogenous flTDP43 protein levels upon EGFP-TDP43 overexpression, consistent with modest autoregulation (**Figure 2c**). We previously developed customized antibodies^19^ that recognize the C-terminal 18-amino acids unique to sTDP43; this segment is absent from flTDP43, permitting selective detection of sTDP43 by immunoblotting. In N2a cells expressing EGFP-TDP43 and treated with SMG1i, we again observed an increase in the 33kDa band corresponding to endogenous sTDP43, providing additional evidence of sTDP43 protein accumulation in response to TDP43 autoregulation and NMD inhibition (**Supplemental Figure 1c**-**d**).

**Figure 2:**
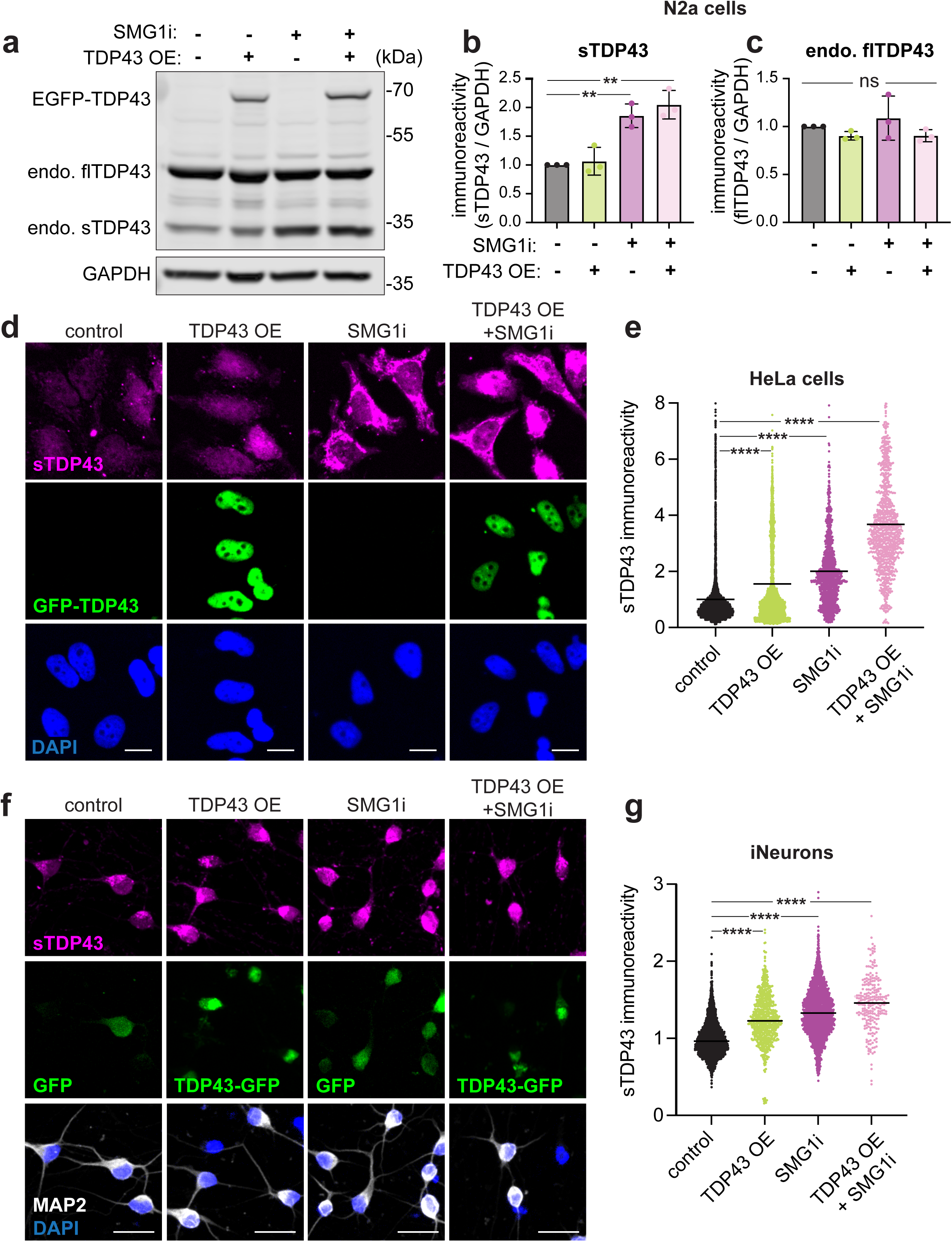
sTARDBP transcripts that escape NMD undergo translation. (**a**) Western blot of N2a cells overexpressing GFP-TDP43 and/or treated with SMG1i, probed with an N-terminal TDP43 antibody that detects all TDP43 species of interest. (**b**) Quantification of bands corresponding to sTDP43 in (**a**), showing increase in sTDP43 with TDP43 OE and SMG1i. (**c**) Quantification of endogenous full-length (endo. fl)TDP43 immunoreactivity in (**a**), demonstrating no significant change following TDP43 OE or NMD inhibition. (**d**) Immunostaining for sTDP43 in Flp-In GFP-TDP43 HeLa cells overexpressing GFP-TDP43 and/or treated with SMG1i. Scale bars: 50µm. (**e**) Quantification of sTDP43 immunoreactivity in (**d**), showing synergistic increase in sTDP43 with GFP-TDP43 OE and NMD inhibition. N=number of cells analyzed: control n=5774, TDP43 OE n=2996, SMG1i n=1245, TDP43+SMG1i n=975. (**f**) Immunostaining for sTDP43 in iNeurons transduced with GFP or TDP43-GFP, and treated with vehicle or SMG1i for 48h. Scale bars: 25µm (**g**) Quantification of sTDP43 immunoreactivity in (**f**). Control n=2874, TDP43 OE n=720, SMG1i n=2138, TDP43+SMG1i n=229. (**b**-**g**) Data combined from 3 replicates. **p<0.01, ****p<0.0001 by 2-way ANOVA with Tukey’s multiple comparisons test.

We also confirmed these findings by immunofluorescence. Flp-In GFP-TDP43 HeLa cells^41^ were treated with DOX (TDP43 OE) and/or SMG1i for 48h, then immunostained for endogenous sTDP43 before imaging the cells by fluorescence microscopy (**Figure 2d**, **e**). In keeping with our observations and those of others^19–20^, sTDP43 was localized primarily within the cytosol, contrasting with the distribution of GFP-TDP43. Once more, we noted significant increases in sTDP43 abundance upon TDP43 OE and SMG1i treatment. Moreover, combined TDP43 OE and SMG1i application synergistically boosted sTDP43 immunoreactivity (**Figure 2e**), analogous to what we observed at the RNA level (**Figure 1**). Similar results were obtained in human iNeurons overexpressing TDP43-EGFP and/or treated with SMG1i (**Figure 2f**, **g**): while TDP43 autoregulation and NMD inhibition increased sTDP43 protein levels on their own, combining both treatments led to a substantial increase in endogenous sTDP43 production.

Antibodies may display off-target or unpredictable reactivity, particularly for low-abundance proteins such as sTDP43. We therefore developed two distinct, antibody-independent approaches to confirm the effects of TDP43 autoregulation and NMD on sTDP43 levels. First, we designed and created a minigene reporter of sTDP43 by inserting the EGFP open reading frame (ORF) upstream of *TARDBP* exon 6 and the proximal 3’UTR, sequences that include the sTDP43-specific splice donor and splice acceptor sites, respectively. To track sTDP43-specific splicing, an mApple ORF was inserted immediately downstream of the sTDP43 stop codon in exon 7. In cells expressing the reporter, the ratio of mApple (RFP) / EGFP (GFP) fluorescence represents a direct readout of sTDP43-productive splicing (**Figure 3a**). We co-expressed this splicing reporter with vectors encoding a far-red fluorescent protein (iRFP, a control) or TDP43 fused to iRFP (TDP43 OE) in rat primary neurons, then measured changes in the ratio of RFP/GFP fluorescence for each neuron over time via automated fluorescence microscopy. Consistent with TDP43 autoregulation inducing sTDP43-productive splicing, we detected a significant increase in the single-cell RFP/GFP ratio over time in neurons overexpressing TDP43-iRFP, compared to iRFP alone (**Figure 3b**).

**Figure 3:**
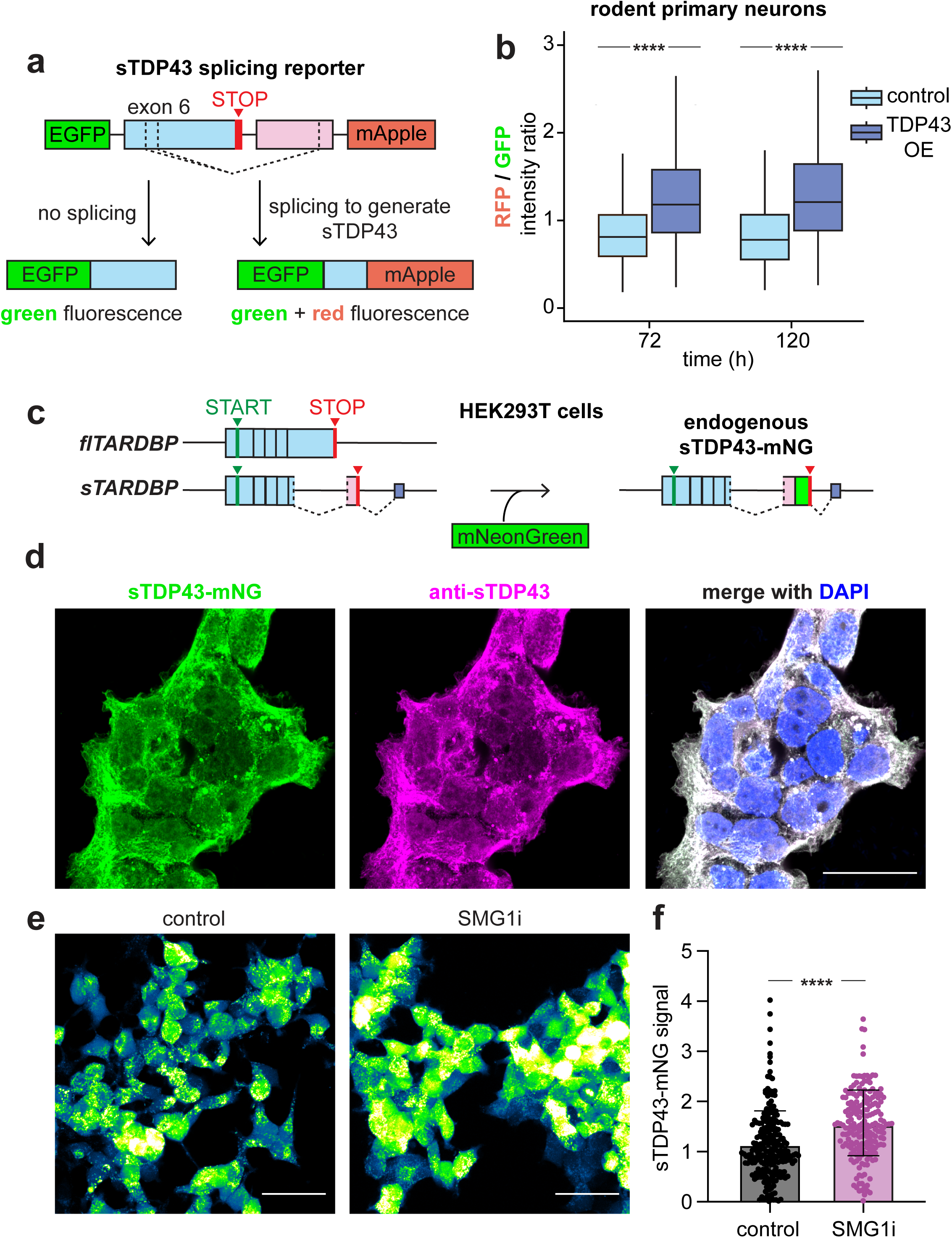
Antibody-independent approaches confirm strict control of sTDP43 production by TDP43 autoregulation and NMD. (**a**) Schematic of sTDP43 splicing reporter. EGFP is fused upstream of *TARDBP* exon 6 (blue) encoding *sTARDBP-1* and *sTARDBP-2* splice donors. mApple is fused downstream of termination codon, the *sTARDBP* splice acceptor site and polyadenylation signal within the *TARDBP* 3’UTR (pink). At baseline, the reporter fluoresces green only, while sTDP43-specific splicing drives green and red fluorescence. (**b**) TDP43 OE significantly increases the RFP / GFP ratio, indicative of an increase in sTDP43-specific splicing, in rodent mixed primary cortical neurons. 72h control n=719, 72h TDP43 OE n=406, 120h control n=455, 120h TDP43 OE n=195. ****p<0.0001 by Wilcox signed-rank test combined and stratified among 6 biological replicates. (**c**) Schematic of mNG insertion and labeling of endogenous sTDP43 in HEK293T cells. (**d**) Intrinsic sTDP43-mNG signal overlaps with immunoreactivity from a sTDP43-specific antibody. Scale bars: 25µm. (**e**) sTDP43-mNG fluorescence in live cells treated with vehicle or SMG1i for 24h. Scale bars: 50µm. (**f**) SMG1i significantly increases intrinsic sTDP43-mNG signal. N=number of cells analyzed among 3 biological replicates: control n=221, SMG1i n=189. ****p<0.0001 by unpaired t-test.

The sTDP43 splicing reporter lacks the entirety of the *TARDBP* 3’UTR, and is therefore missing key elements that would target it for destruction by NMD (i.e. additional splicing events >50nt downstream of the stop codon)^55^. To examine NMD-dependent sTDP43 regulation, we instead utilized the native genetic context by inserting an open reading frame encoding the ultrabright fluorescent protein mNeonGreen (mNG) just upstream of the *sTARDBP* termination codon (**Figure 3c**). Sanger sequencing confirmed the insertion of mNG at the correct location. The endogenous mNG signal overlapped closely with the cytosolic, punctate distribution of sTDP43 as detected by immunostaining using three separate sTDP43-selective antibodies (**Figure 3d**, **Supplemental Figure 1e**). In contrast, sTDP43-mNG puncta showed little direct overlap with cytosolic organelles such as lysosomes, stress granules and processing (P)-bodies under normal conditions (**Supplemental Figure 2**). Upon addition of sodium arsenite, an oxidative stressor that induces stress granule formation and P-body enlargement^56–57^, sTDP43-mNG appeared to partially overlap with the periphery of these structures. To validate our previous findings, we added SMG1i to the sTDP43-mNG HEK293T cells for 24h and imaged the intrinsic mNG signal in live cells. Confocal fluorescence microscopy confirmed a significant mNG signal increase in SMG1i-treated cells compared to vehicle control (**Figure 3e**, **f**). Together, these data confirm that sTDP43-encoding transcripts are created through TDP43 autoregulation and subject to NMD.

### sTDP43 protein is targeted for rapid degradation

We next asked whether sTDP43 levels are regulated not just at the RNA level, but also post-translationally through protein turnover mechanisms. To do so, we turned to optical pulse labeling (OPL), a non-invasive method for measuring protein turnover *in situ*^48,52,58–61^. OPL takes advantage of dendra2, a photoconvertible protein that fluoresces green at baseline, but upon exposure to UV light (405nm), irreversibly converts to a red fluorophore^62^. By measuring the decay of the red signal over time in individual cells, we calculated a half-life for dendra2-fused proteins. In previous studies, the turnover of flTDP43 and flTDP43-dendra2—as measured by metabolic pulse-chase and OPL—were nearly identical to one another^59^, confirming the validity of OPL for tracking TDP43 clearance in cultured neurons.

To measure sTDP43 half-life, we transfected rodent primary cortical neurons with plasmids encoding dendra2 or C-terminal fusions of sTDP43-dendra2 and flTDP43-dendra2. All neurons were co-transfected with a plasmid encoding EGFP to serve as a cell marker. Using automated fluorescence microscopy, we imaged transfected neurons regularly before and after photoconversion with UV light (**Figure 4a**, **b**). As expected, based on previous studies, flTDP43-dendra2 half-life was ∼46h^59,61^. In contrast, the measured half-life of sTDP43-dendra2 was significantly shorter (∼18h; **Figure 4c**) suggesting that the truncated protein is rapidly cleared compared to flTDP43-dendra2.

**Figure 4.**
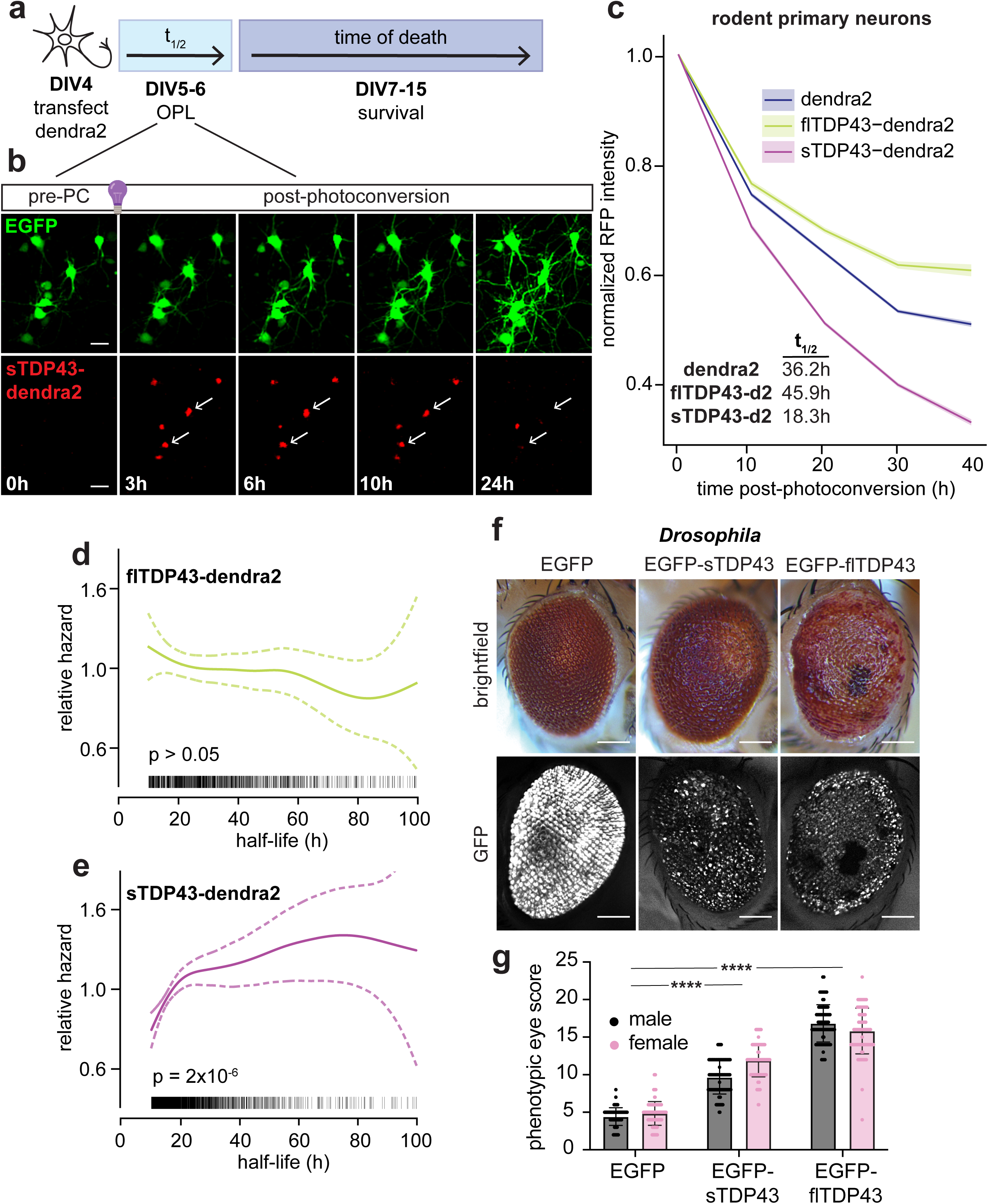
sTDP43 protein is targeted for rapid degradation. (**a**) Schematic of optical pulse labeling (OPL) and survival experiments. (**b**) Representative images of transfected neurons pre- and post-photoconversion (PC). Scale bars: 50µm. (**c**) Decay of red fluorescent signal for dendra2, sTDP43-dendra2 (d2) and flTDP43-dendra2 (d2), normalized to the initial measurement post-photoconversion. Data combined among 16 biological replicates. (**d**, **e**) Cox penalized spline models illustrating the relationship between single-cell flTDP43-d2 (**d**) or sTDP43-d2 (**e**) half-life and the relative risk of death as judged by automated fluorescence microscopy and survival analysis. Tick marks along the x-axis indicate half-life measurements from individual neurons. Dotted lines, 95% CI. N=number of neurons among 4 biological replicates, flTDP43-dendra2 n=1041, sTDP43-dendra2 n=1857. p-value determined by likelihood ratio test. (**f**) Representative images and (**g**) quantification of rough eye phenotype in transgenic *Drosophila* overexpressing EGFP, EGFP-sTDP43 or EGFP-flTDP43 via the gmr-Gal4 driver. Both EGFP-flTDP43 and EGFP-sTDP43 are significantly more toxic than EGFP alone. ****p<0.0001 by 2-way ANOVA with Tukey’s. Scale bars: 100µm.

Our earlier work indicated that flTDP43 is degraded via the ubiquitin-proteasome system (UPS) as well as macroautophagy^48,59,61^. To determine if sTDP43 turnover is accomplished through similar mechanisms, we performed OPL for sTDP43-dendra2 in the presence of pharmacological inhibitors of these pathways (**Supplemental Figure 3a**-**f**). We found that both MG132 (**Supplemental Figure 3a**) and Bafilomycin-A1 (BafA1), (**Supplemental Figure 3d**), inhibitors of the UPS and autophagy, respectively, significantly extended the half-life of sTDP43-dendra2. Together, these findings indicate that sTDP43 is an unstable protein that is degraded through a combination of the UPS and macroautophagy, further contributing to its low steady-state levels.

Individual neurons vary widely in their ability to degrade sTDP43-dendra2, with half-life values ranging from 10h to over 200h for each neuron. Such variability at the single-cell level is in keeping with what we have previously observed for flTDP43-dendra2 and reporters of macroautophagy (i.e. LC3-dendra2)^52,59^. We reasoned that if sTDP43 accumulation is toxic, then the survival of individual neurons should be proportional to each cell’s ability to degrade sTDP43. To address this, we first measured the turnover of sTDP43-dendra2 and flTDP43-dendra2 24-48h after transfection of rodent primary neurons, then tracked the survival of individual cells for 8d using automated fluorescence microscopy^19,49,61,63–64^ (**Fig. 4a**). To determine if protein half-life measured at the outset of the experiment is an accurate predictor of survival, we applied Cox proportional hazards analysis, and visualized the relationship between single-cell half-life and the relative risk of death through a penalized spline model^59,65^. Although there was no clear effect of flTDP43-dendra2 half-life on survival (**Figure 4d**), we observed a striking and direct relation between sTDP43-dendra2 turnover and survival (**Figure 4e**) —the relative risk was 50% for neurons displaying shorter sTDP43 half-life (<30h), while toxicity was ∼50% higher for cells exhibiting a longer sTDP43 half-life (>30h). These data imply that sTDP43 stabilization is strongly associated with an increased risk of cell death.

To explore this possibility in more detail, and examine the consequences of sTDP43 accumulation *in vivo*, we utilized a sTDP43 cDNA construct that avoids degradation by NMD, and whose expression is high enough to compensate for post-translational destabilization of the protein. Using the yeast Gal4-UAS system and a gmr-Gal4 driver, we overexpressed EGFP, EGFP-sTDP43 or EGFP-flTDP43 in the *Drosophila* eye. After confirming transgene expression via fluorescence microscopy (**Figure 4f**), we assessed rough eye phenotypes in each condition. Similar to prior studies, EGFP-flTDP43 overexpression led to severe damage^66–71^, as indicated by necrotic lesions and collapse of the convex surface of the eye (**Figure 4f**, **g**). sTDP43 overexpression, on the other hand, resulted in significant but milder toxicity in comparison to EGFP-flTDP43. Despite a trend towards lower levels for EGFP-sTDP43, no significant differences in transgene expression at the RNA or protein levels were detected by qRT-PCR or immunoblotting, respectively, for EGFP-sTDP43 and EGFP-flTDP43 (**Supplemental Figure 3g**-**i**). In keeping with prior studies in rodent primary neurons^19^, these data confirm that sTDP43 accumulation is neurotoxic both *in cellulo* and *in vivo*.

### sTDP43-mediated toxicity requires functional TDP43 dimerization and RNA binding

We next sought to understand the mechanism by which sTDP43 drives toxicity. Our previous investigations demonstrated not just sTDP43 deposition within cytosolic inclusions (i.e. gain-of-function), but also nuclear exclusion and cytosolic retention of endogenous flTDP43 by sTDP43 (i.e. loss-of-function)^19^. Given the critical function of the TDP43 N-terminus in mediating TDP43 oligomerization^72–73^, and our prior data showing that sTDP43 interacts with flTDP43^19^, we suspected that sTDP43 may be sequestering endogenous flTDP43 via its intact N-terminus. To test this, we mutated four residues within the N-terminus of sTDP43 that are essential for TDP43 oligomerization (E17/21A, R52/55A, “4M”)^73^. We also created a phosphomimetic substitution (S48E) predicted to interrupt TDP43 oligomerization^74^ (**Figure 5a**, **b**). To validate the oligomerization deficits induced by these mutations, we created HaloTag (Halo) fusions of sTDP43, sTDP43(4M) and sTDP43(S48E). Each was expressed in HEK293T cells, followed by HaloLink immunoaffinity purification and immunoblotting for endogenous flTDP43 pulled down by each variant (**Supplemental Figure 4a**). While sTDP43-Halo effectively pulled down endogenous flTDP43 (endo.flTDP43), this interaction was minimized by the 4M mutations within the sTDP43 N-terminus. These mutations also prevented the interaction of sTDP43 with itself (sTDP43-Halo), suggesting that sTDP43 is capable of homooligomerization as well as heterooligomerization with endogenous flTDP43. In contrast, the S48E substitution had a more subtle effect on the ability of sTDP43 to interact with endogenous flTDP43 or itself (**Supplemental Figure 4b**).

**Figure 5.**
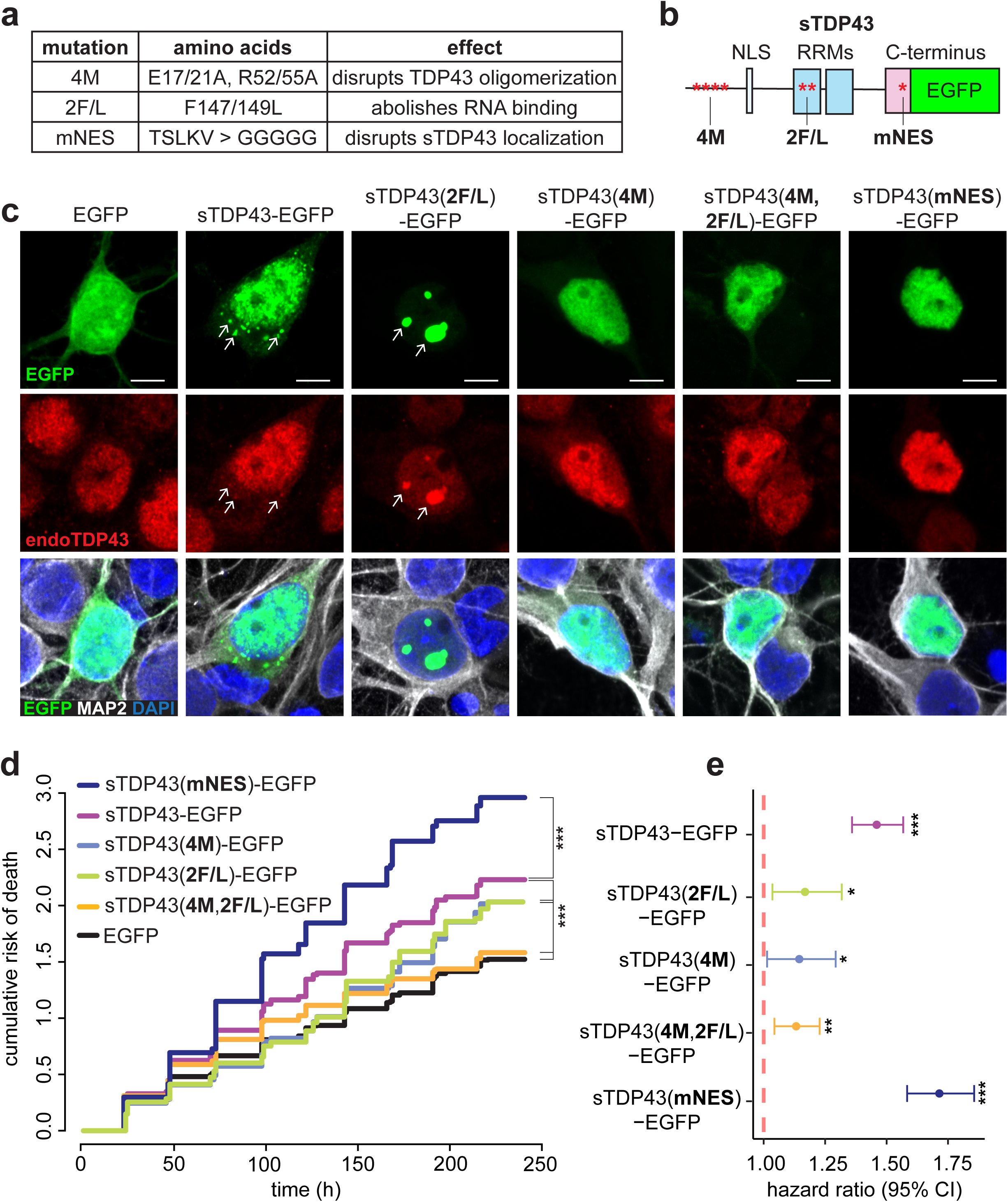
sTDP43-mediated toxicity requires functional TDP43 dimerization and RNA binding. (**a**-**b**) Descriptions and location of engineered sTDP43 mutations. (**c**) Representative images of EGFP-tagged sTDP43 variants in rodent primary neurons. Scale bars: 5µm. (**d**) Cumulative hazard and (**e**) forest plot of hazard ratios of sTDP43 variants, determined by automated microscopy. The negative control (EGFP) serves as a reference (Hazard ratio=1). N=number of neurons, combined and stratified among 6 biological replicates: EGFP n=2047, sTDP43-EGFP n=1641, sTDP43(4M)-EGFP n=469, sTDP43(2F/L)-EGFP n=484, sTDP43(4M,2F/L)-EGFP n=1335, sTDP43(mNES)-EGFP n=1263. *p<0.05, **p<0.01, ***p<0.001 by Cox proportional hazard analysis.

Our prior investigations indicated that gain-of-function toxicity from flTDP43 overexpression relies upon RNA binding and the intramolecular interactions between RNA recognition motifs (RRMs) 1 and 2^61^. Given the intact RRMs within sTDP43, we surmised that sTDP43 may also be able to bind RNA and elicit analogous gain-of-function toxicity when overexpressed. To address this, we again expressed sTDP43-Halo in HEK293T cells, and purified sTDP43-Halo using HaloTrap after UV crosslinking (**Supplemental Figure 4a**). While sTDP43-Halo pulled down canonical TDP43-bound RNAs such as *YTHDF1* and *ACTIN*, the introduction of a double F147L/F149L (2F/L) mutation in RRM1—sufficient to block RNA binding by flTDP43 in previous studies^61,75–76^—completely inhibited recognition of these substrates by sTDP43 (**Supplemental Figure 4c**).

We also examined how disrupting sTDP43 oligomerization and RNA binding impacted its localization. In HEK293T cells, sTDP43-EGFP accumulated within cytosolic puncta (**Supplemental Figure 5**). In contrast, both sTDP43(4M)-EGFP and sTDP43(2F/L)-EGFP were primarily nuclear, suggesting N-terminal interactions and RNA binding are important for cytosolic sTDP43 distribution. The large nuclear droplets formed by sTDP43(2F/L)-EGFP were eliminated by the 4M mutation, highlighting the importance of N-terminal interactions for intranuclear phase separation. As a control, we also expressed sTDP43(mNES)-EGFP, a variant of sTDP43 carrying a mutated version of the predicted NES within the unique C-terminus of sTDP43^19^. This construct displayed nuclear localization similar to that of sTDP43(4M)-EGFP and sTDP43(2F/L)-EGFP. Analogous results were observed in rodent primary cortical neurons expressing each of the sTDP43-EGFP constructs, as well as identical variants in flTDP43-EGFP (**Figure 5c**, **Supplemental Figure 6a**, **b**). In these neurons, we also tracked the sequestration of endogenous flTDP43 by exogenous sTDP43-EGFP variants. As expected, flTDP43-EGFP was nuclear, while sTDP43-EGFP accumulated within cytoplasmic puncta that colocalized with endogenous flTDP43. The nuclear droplets formed by RNA binding-deficient sTDP43(2F/L)-EGFP also effectively sequestered endogenous flTDP43. Both the 4M and mNES mutations resulted in nuclear redistribution of sTDP43-EGFP, as in HEK293T cells, but given the nuclear predominance of flTDP43, it was difficult to determine if there was any true colocalization of sTDP43(4M)-EGFP or sTDP43(mNES)-EGFP with endogenous flTDP43 by this technique.

To discriminate whether RNA binding or N-terminal interactions with endogenous flTDP43 are relevant for sTDP43-related toxicity, we again turned to automated fluorescence microscopy and survival analysis^19,49,61,63–64^. Here, rodent primary cortical neurons were co-transfected with plasmids encoding sTDP43-EGFP variants as well as a separate vector encoding a red fluorescent cell marker, mApple. We then imaged transfected cells at regular 24h intervals over the course of 10d and compared survival between conditions by Cox proportional hazards analysis.

In harmony with our earlier studies^19^ and data from *Drosophila* models, sTDP43-EGFP overexpression led to significant toxicity in comparison with EGFP alone (**Figure 5d**, **e**). Variants carrying the 2F/L and 4M mutations displayed marked reductions in risk of death compared to wild-type (WT) sTDP43-EGFP, indicating that both RNA binding and N-terminal interactions drive sTDP43-dependent toxicity. The double 4M,2F/L mutant exhibited an additive benefit, suggesting that RNA binding and N-terminal interactions enhance toxicity through independent pathways. In parallel experiments, we observed a similar pattern for flTDP43-EGFP: toxicity due to this construct was dependent not just on RNA binding, as noted previously^61^, but also N-terminal interactions (**Supplemental Figure 6c**, **d**). Together, these results demonstrate that neurotoxicity driven by sTDP43 requires functional RNA binding as well as N-terminal interactions.

Because the 4M and 2F/L mutations both resulted in nuclear retention of sTDP43, we questioned whether the reduction of toxicity observed in association with these mutations might be secondary to changes in localization. Primary neurons were transfected with sTDP43(mNES)-EGFP, which likewise demonstrates nuclear localization, and the survival of these cells was tracked by automated microscopy. In contrast to the 4M and 2F/L mutations, disruption of the predicted NES led to a striking increase in toxicity for sTDP43(mNES)-EGFP in comparison to sTDP43(WT)-EGFP (**Figure 5d, e**). These results suggest that cytoplasmic localization is not necessary for toxicity; rather, sTDP43 cytosolic deposition may partially mitigate toxicity by limiting heterooligomerization with flTDP43 or access to nuclear RNA.

Due to the stochastic nature of transient transfection, the number of cells included within each condition was inconsistent. To uncover any bias in the results arising from this imbalance, we bootstrapped the data by replotting the survival of 100 randomly selected cells from each experiment, for 10 iterations. Cumulative hazard plots and hazard ratios for both sTDP43 (**Supplemental Figure 7a**) and flTDP43 (**Supplemental Figure 7b**) survival data showed no significant differences relative to original analyses (**Figure 5d**, **e** and **Supplemental Figure 6c**, **d**), arguing against skew from uneven population sizes.

### sTDP43 accumulation drives loss of TDP43 function

To determine if sTDP43 does indeed oligomerize with nuclear flTDP43, we used disuccinimidyl glutarate (DSG), a membrane permeable cross-linker, in HEK293T cells expressing each EGFP-tagged sTDP43 variant (**Supplemental Figure 8a**, **b**) to analyze protein-protein interactions. We detected a prominent shift towards high molecular weight oligomers after DSG application in cells expressing sTDP43-EGFP, indicating successful oligomerization of sTDP43 with itself and/or endogenous flTDP43. These data are consistent with the ability of sTDP43 to bind with—and potentially impact the function of— flTDP43. As expected, oligomerization-deficient sTDP43 variants (sTDP43(4M)-EGFP and sTDP43(4M,2F/L)-EGFP) displayed significantly less oligomerization than WT sTDP43-EGFP, but the 2F/L mutations alone did not affect oligomerization. Despite the nuclear localization of sTDP43(mNES)-EGFP, this variant exhibited significantly less oligomerization than WT sTDP43-EGFP, consistent with previous studies indicating the importance of an intact sTDP43 C-terminus for oligomerization and aggregation^77^. Exogenous sTDP43(mNES)-EGFP may therefore elicit toxicity via RNA binding, rather than oligomerization with flTDP43.

To investigate this further, we next assessed the splicing capacity of endogenous flTDP43 in the presence of each engineered sTDP43 variant using a minigene reporter encoding of the cystic fibrosis transmembrane conductance regulator (CFTR) exon 9. Under normal conditions, TDP43 effectively blocks the inclusion of this exon^75,78^, generating an increase in the fully spliced product. However, when TDP43 function is compromised, both the unspliced product and an intermediate ‘cryptic’ splice product are apparent^61^ (**Supplemental Figure 8c**). As a control for TDP43 gain-of-function, we overexpressed flTDP43-EGFP alongside the CFTR minigene; as expected, this led to a significant increase in the ratio of spliced / unspliced product compared to EGFP alone (**Supplemental Figure 8d, e**). We also expressed RNA-binding deficient flTDP43(2F/L)-EGFP, which acts in a dominant-negative manner by sequestering endogenous flTDP43 within nuclear droplets^61^ (**Supplemental Figure 6a**, **b**), resulting in a significant decrease in the ratio of spliced / unspliced product. WT sTDP43-EGFP mimicked this result, eliciting a reduction in the ratio of spliced / unspliced product compared to that produced by EGFP, and indicating a dominant-negative effect of sTDP43 on native flTDP43 splicing activity. The RNA binding-deficient sTDP43 variant (sTDP43(2F/L)-EGFP) also caused a loss of TDP43 function; however, the oligomerization-deficient sTDP43 (sTDP43(4M)-EGFP) and double mutant (sTDP43(4M,2F/L)-EGFP) displayed no significant difference in CFTR splicing compared to EGFP. Conversely, expression of sTDP43(mNES)-EGFP led to cryptic splicing of the CFTR minigene (**Supplemental Figure 8d**, **f**), but not exon 9 inclusion as seen with flTDP43 loss of function, in keeping with its relatively inefficient oligomerization (**Supplemental Figure 8a**, **b**). Collectively, these observations imply that sTDP43 interferes with the function of endogenous flTDP43 via its N-terminal oligomerization and RNA binding properties.

One of the most consistent missplicing events associated with TDP43 pathology in ALS is the downregulation of *Stathmin-2* (*STMN2*), caused by the inclusion of a cryptic exon (2A) in the mature *STMN2* transcript upon TDP43 loss of function^8–10,13^. To determine if sTDP43 is capable of blocking flTDP43 activity and reproducing *STMN2* missplicing, we created a minigene reporter that includes *STMN2* exons 1 and 2, separated by a truncated intron encoding mNG in frame with cryptic exon 2A. Under normal conditions, mNG should be spliced out of the mature transcript. Upon loss of functional TDP43, however, cryptic exon 2A and mNG will be spliced in, producing a measurable increase in green fluorescence (**Figure 6a**). We expressed this reporter in human neuroblastoma (M17) cells together with non-targeting (NT) shRNA or shRNA directed against *TARDBP* as negative and positive controls, respectively. We then imaged the cells using automated fluorescence microscopy 24h after transfection. As expected, we detected a significant increase in mNG intensity in M17 cells expressing the *STMN2* reporter and *TARDBP* shRNA, in comparison to NT shRNA. Co-expression with sTDP43 elicited a similar increase in mNG fluorescence, relative to a control vector expressing iRFP, consistent with the negative influence of sTDP43 on the splicing activity of flTDP43 (**Figure 6b**, **c**).

**Figure 6.**
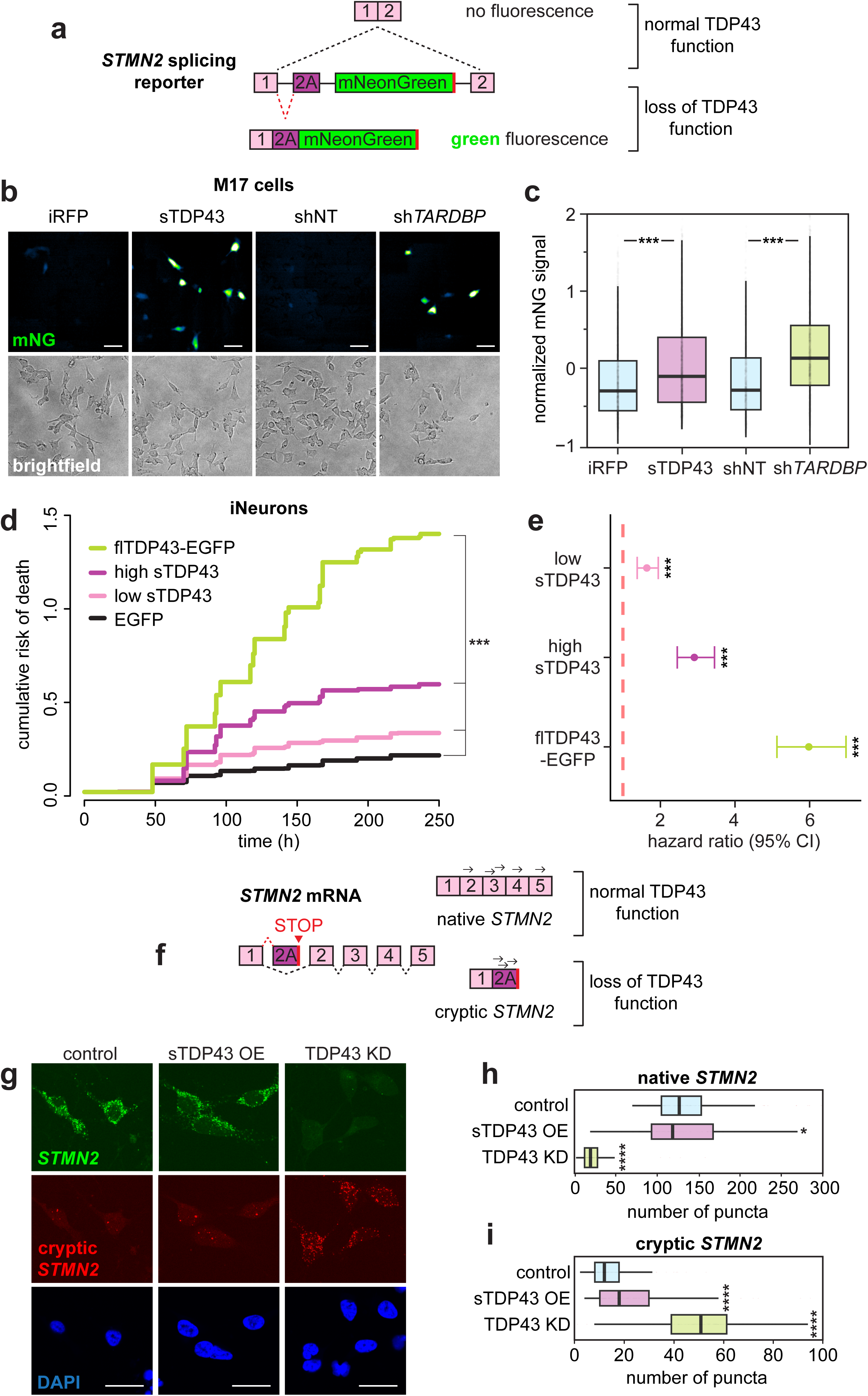
sTDP43 accumulation drives loss of TDP43 function. (**a**) Schematic of *STMN2* splicing reporter. Upon loss of TDP43 function, cryptic exon 2A is included in the mature *STMN2* transcript as detected by mNG (mNeonGreen) fluorescence. (**b**) Representative images and (**c**) quantification of *STMN2* reporter green fluorescence signal in M17 cells transfected with iRFP, sTDP43, shNT, or sh*TARDBP*. sTDP43 overexpression recapitulates the increase in mNG signal detected upon TDP43 knockdown. ***p<0.0001 by 1-way ANOVA with Tukey’s. Scale bars: 100µm (**d**) Cumulative hazard and (**e**) forest plot of iNeurons transduced with EGFP, flTDP43-EGFP and high and low titers untagged sTDP43 virus. N=number of neurons, combined and stratified among 3 biological replicates: EGFP n=804, low sTDP43 n=702, high sTDP43 n=527, flTDP43-EGFP n=681. ***p<0.001 by Cox proportional hazard analysis. (**f**) Schematic of endogenous *STMN2* splicing. Arrows indicate probes used to amplify native and cryptic transcripts. (**g**) Representative images and (**h**-**i**) quantification of HCR FISH for (**h**) native and (**i**) cryptic exon-containing *Stathmin-2 (STMN2)* transcripts in iNeurons upon sTDP43 overexpression (OE) and TDP43 knockdown (KD). *p<0.05, ****p<0.0001 by 1-way ANOVA with Tukey’s. Scale bars: 25µm.

Based on these data, we next investigated whether sTDP43 accumulation causes toxicity and missplicing of endogenous disease-associated transcripts in human neurons. iNeurons were transduced with lentiviral vectors expressing untagged sTDP43, and the cells were followed for 10d in culture through automated longitudinal microscopy. As a positive control, we overexpressed flTDP43 fused with EGFP, which induces dose-dependent toxicity when expressed in neurons^22,61,63^. (**Figure 6d**, **e**). The time of death for individual neurons was determined by manually assisted survival analysis, using the same set of criteria (cell rounding, retraction of processes) utilized for automated analysis. Consistent with what we observed in *Drosophila* (**Figure 4f**) and rodent primary neurons (**Figure 5**), these studies demonstrated dose-dependent neurotoxicity with sTDP43 overexpression, albeit modest compared to flTDP43 overexpression. To determine whether sTDP43 overexpression likewise causes TDP43-dependent missplicing in neurons, we performed hybridization chain reaction fluorescent *in situ* hybridization (HCR FISH) for *STMN2*, a transcript uniquely expressed and spliced by TDP43 in human neurons^8–9^. Loss of TDP43 function leads to the inclusion of a cryptic exon (2A) and truncation of the *STMN2* transcript, together with a reduction in full-length *STMN2* mRNA (**Figure 6f**). In unmodified iNeurons, full-length (native) *STMN2* is the dominant isoform, as expected in healthy cells containing physiological levels of TDP43 (**Figure 6g**-**i**). However, upon TDP43 depletion, native *STMN2* levels drop dramatically, accompanied by a corresponding increase in the cryptic *STMN2* product (**Figure 6g**-**i**). iNeurons transduced with sTDP43 lentivirus demonstrate a subtle drop in full-length *STMN2* and a modest but significant increase in cryptic *STMN2*, confirming that sTDP43 interferes with flTDP43 splicing activity in human neurons.

## Discussion

In this study, we show that sTDP43, an alternatively spliced, shortened variant of TDP43, is generated as a byproduct of TDP43 autoregulation and governed tightly by NMD. sTDP43 transcripts that escape NMD and undergo translation are rapidly cleared by the UPS and macroautophagy, further contributing to low steady-state levels. Bypassing these regulatory mechanisms by overexpressing sTDP43 in primary rodent neurons drives toxicity that is dependent upon the ability of sTDP43 to bind RNA as well as N-terminal oligomerization with flTDP43. This suggests both gain- and loss-of-function disease mechanisms. We observed analogous toxicity in *Drosophila* eye and human iPSC-derived neurons expressing sTDP43, together with a loss of endogenous flTDP43 splicing activity. These results uncover a series of regulatory pathways for maintaining low sTDP43 abundance, providing a potential explanation for the relatively scarcity of sTDP43 and the inability to detect these isoforms in prior studies. Our findings also imply that sTDP43 accumulation in ALS, which we previously observed^19^, may interfere with flTDP43 activity in a dominant-negative manner, thereby recapitulating the functional loss of TDP43 activity characteristic of ALS, FTLD-TDP and other TDP43 proteinopathies.

TDP43 regulates its own expression through an intricate negative feedback loop in which TDP43 binds to its own mRNA transcript, triggering alternative splicing within exon 6 and the 3’UTR^4,18,38^. Previous studies hinted at the presence of truncated variants produced via TDP43 autoregulation that are targeted by NMD^4,40^. Here, we connect these events to sTDP43 splice variants identified in hyperexcitable neurons and ALS patient tissue^19–20^. It is unclear whether intrinsic deficiencies in NMD may contribute to the accumulation of sTDP43 in ALS or FTLD; although initial studies suggested inefficient NMD in ALS/FTLD models^63,79–80^, more recent investigations argue against this hypothesis^81^. Similarly, the lack of nuclear TDP43 in affected neurons in ALS/FTLD-TDP is associated with reduced autoregulation^40,82^, predicting decreases rather than the observed increase in sTDP43 abundance. We propose that disease-associated sTDP43 accumulation may be secondary to inefficient post-translational degradation of the protein. In support of this, autophagic flux declines with advanced age^83^, and age remains the strongest risk factor for ALS, FTLD-TDP and related neurodegenerative conditions. Additionally, pathogenic mutations in *C9ORF72*, *VCP*, *UBQLN2*, *GRN*, *SQSTM1*, and *TBK1* all impair autophagy and result in ALS/FTLD-TDP with TDP43 pathology^84–90^. As flTDP43 is also a substrate of macroautophagy^59,61^, and this pathway is primarily active within the cytosol, deficiencies in this pathway may lead to the cytosolic deposition of both flTDP43 and sTDP43. Together, these observations suggest a potential means by which age-related or genetic impairments in autophagy result in key neuropathological features of disease, including cytosolic aggregates of sTDP43/flTDP43 and loss of nuclear flTDP43 splicing activity.

In previous work, we determined that sTDP43 variants are upregulated by neuronal hyperexcitability^19^, a consistent early feature of ALS and FTLD-TDP^91–96^. Although the precise relation between neuronal activity, TDP43 autoregulation, and NMD remains unknown, our investigations suggest pivotal connections among these pathways that may be crucial for disease pathogenesis. Some of the best-known examples of activity-dependent genes, including *ARC* (activity-regulated cytoskeleton-associated protein), are NMD substrates whose abundance rapidly increases upon neuronal stimulation^97^. Neuronal hyperactivity also upregulates the expression of hnRNPA2B1— an RNA binding protein related to TDP43 and is itself associated with familial ALS^98^— through an NMD-dependent mechanism^99^. One possibility is that neuronal hyperexcitability, as seen in early stages of ALS and FTLD-TDP^100–101^, leads to sTDP43 accumulation secondary to persistent downregulation of NMD. Confirmation of this hypothesis, as well as the potential mechanisms responsible for activity dependent NMD impairment, will require additional investigation.

We detected consistent changes in sTDP43 at the RNA level by qRT-PCR, and at the protein level by immunofluorescence, in response to NMD and TDP43 autoregulation, in several different cell types. By western blot, sTDP43 protein was only detectable in mouse N2a cells; in comparison, we were unable to accurately measure sTDP43 by western blot in human cell lines or human neurons (data not shown). These results are consistent with prior studies the presence of sTDP43 by immunoblotting and mass spectroscopy in mouse, but not human, brain tissue^18^. We and others have previously found sTDP43 to be highly prone to aggregation^16,19^, suggesting that sTDP43 forms insoluble inclusions with buried epitopes upon its induction. These observations also imply unique behavior of sTDP43 within rodents and humans, despite the high degree of sTDP43 sequence conservation among species. Even so, we noted significant changes in sTDP43 levels in human cells using immunofluorescence, highlighting the variability in sTDP43 abundance between cells, and the sensitivity of single-cell approaches over population-based measures such as western blotting.

To bypass any issues that may arise from buried epitopes and the resulting failure to detect endogenous sTDP43, we created knock-in HEK293T cells in which endogenous sTDP43 is tagged with mNG. In these cells, we found minimal colocalization between endogenous sTDP43-mNG and a series of cytosolic organelles at baseline. Following treatment with sodium arsenite, however, sTDP43-mNG appears to associate with both stress granules (marked by G3BP1) and processing (P)-bodies (marked by Xrn1), two membraneless organelles involved in RNA processing. These results are consistent with previous observations showing overexpressed sTDP43 co-purifies and colocalizes with stress granule and P-body components^16–17,19^. While both organelles are associated with translational stalling and RNA stabilization under conditions of cellular stress, P-bodies are uniquely involved in RNA decapping and degradation^102–103^. This, together with findings that sTDP43 retains the ability to bind RNA, may suggest a potential function for sTDP43 in the maintenance of cytosolic RNA stability. Because sTDP43 lacks the C-terminal domain of flTDP43 that is required for many protein-protein interactions, we suspect that sTDP43 acts primarily as a transport protein, binding to RNAs and escorting them to stress granules and P-bodies under stress. These and other possible functions for sTDP43 will be pursued in future studies.

Using a series of engineered sTDP43 mutants, we find that sTDP43 accumulation inhibits flTDP43 function through two mechanisms. First, the sTDP43 N-terminal region, which is responsible for oligomerization^73^, enables sTDP43 to heteroligomerize with flTDP43 and directly interrupt its activity. Consistent with this, we previously found that sTDP43 binds and sequesters flTDP43 within cytosolic deposits, akin to its distribution in disease^19^. Here, we show that this phenomenon is associated with disrupted flTDP43 splicing activity. Second, we demonstrate that sTDP43 RRMs can bind a subset of TDP43 RNA substrates, and that this RNA binding capacity is required for sTDP43-mediated interruption of flTDP43 splicing, perhaps indirectly via competitive inhibition. Together, these mechanisms contribute to potent downregulation of flTDP43 function by sTDP43.

Using optical pulse labeling, we found sTDP43-dendra2 is cleared more than twice as fast as flTDP43-dendra2 in an autophagy- and UPS-dependent manner. These results suggest sTDP43 is specifically targeted for degradation, similar to what we previously observed for disease-associated TDP43 mutants that display a greater tendency for aggregation^61^. In combination with the strict regulatory mechanisms that control sTDP43 production (NMD, TDP43 negative feedback), such enhanced post-translational clearance ensures low baseline levels of sTDP43. However, transient and short-lived increases in sTDP43 abundance might be seen under limited circumstances, such as in response to TDP43 autoregulation or neuronal stimulation. One possibility is that such ‘pulses’ of sTDP43 production may serve to acutely block TDP43 splicing activity until flTDP43 protein levels return to steady-state equilibrium. As shown here and in previous studies, the half-life of flTDP43 is approximately 2d^61,63^, providing a critical window for sTDP43 to act as a naturally occurring, short-term, dominant-negative inhibitor of TDP43 activity.

If sTDP43 does, in fact, act as an acute-phase regulator of flTDP43 function, it follows that abnormally sustained sTDP43 production would lead to persistent loss of flTDP43 activity. In keeping with this, we demonstrate that sTDP43 accumulation cryptic results in splicing abnormalities characteristic of TDP43 dysfunction, in association with toxicity *in vitro* (in rodent primary neurons and human iNeurons) and *in vivo* (in *Drosophila*). The events responsible for sustained sTDP43 production are unknown and remain the focus of ongoing investigations. Nevertheless, given longstanding evidence of neuronal hyperexcitability in the early stages of ALS^100–101,104^, and the effect of hyperexcitability on *TARDBP* mRNA and sTDP43 in particular^19^, we suspect that loss of inhibition or intrinsically elevated neuronal activity may be responsible for sTDP43 accumulation in disease. This, in turn, could lead to flTDP43 dysfunction and cytosolic mislocalization, two signature pathological events in ALS and FTLD-TDP.

## Acknowledgements

We thank the patients who graciously donated the skin cells used to make iPSCs. We thank Clotilde Lagier-Tourenne at Massachusetts General Hospital for the Flp-In GFP-TDP43 HeLa cells^41^, Zachary Campbell at the University of Texas at Dallas for generously providing 11j, Yuna Ayala at Saint Louis University for sharing the CFTR plasmid^78^, Dr. Henry Paulson and members of his laboratory at the University of Michigan for sharing antibodies, Dr. Peter Todd and Yi-Ju (LuLu) Tseng at the University of Michigan for their help in preparing RNA, Tonya Kopas at the University of Michigan Vector Core for lentivirus preparation, and the University of Michigan Advanced Genomics core for assisting in next-generation RNA sequencing.

This work was supported by National Institutes of Health (R01NS097542, R01NS113943 and 1R56NS128110-01 to SJB; F31NS134123-01 to MMD; F31NS115257 to NBG; P30AG072931 to the University of Michigan Brain Bank and Alzheimer’s Disease Research Center; and R01NS099280 to PKT). Department of Defense (W81XWH2110182 to SJB), the VA (BLRD BX004842 to PKT), the family of Angela Dobson and Lyndon Welch, the A. Alfred Taubman Medical Research Institute, the Danto Family, Ann Arbor Active Against ALS, and the Robert Packard Center for ALS Research.

## Supplemental Materials and Methods

### Culturing and transfecting HEK293T, HeLa cells and N2a cells

Human embryonic kidney (HEK) 293T, HeLa and Neuro-2a (N2a), and M17 cells were cultured in DMEM (Gibco, 11995065), 10% FBS (Gibco, ILT10082147), 1x Glutamax Supplement (Gibco, 35050-061), and 100 units/mL Pen Strep (Gibco, 15140-122) at 37°C in 5% CO_2_. Cells were transfected with Lipofectamine 2000 (Invitrogen, 11668027) according to the manufacturer’s instructions. To induce GFP-TDP43 overexpression in Flp-In HeLa cells^41^, 1µg/mL doxycycline (Sigma, D3447) was added to the media for at least 24h.

### Primary neuron cell culture and transfection

Cortices from embryonic day (E)19-20 Long-Evans rat embryos were dissected and disassociated, and primary neurons were plated at a density of 6x10^5^ cells/mL in 96-well plates or on coverslips. At *in vitro* day (DIV) 4, neurons were transfected with 100ng of a control fluorescent plasmid to mark cell bodies and 100ng of an experimental construct using Lipofectamine 2000 as previously described^19,22,49,59^. Following transfection, cells were placed in either Neurobasal Complete Media (Neurobasal (Gibco 21103-049), 1x B27 Supplement (Gibco, 17504-044), 1x Glutamax, 100 units/mL Pen Strep), NEUMO photostable medium with SOS (Cell Guidance Systems, M07-500), or BrainPhys Imaging Optimized Medium (Stemcell Technologies, 05796) and incubated at 37°C in 5% CO_2_.

### Ethics statement

All vertebrate animal work was approved by the Committee on the Use and Care of Animals (UCUCA) at the University of Michigan (UM), and all experiments were performed in accordance with UCUCA guidelines. Rats (*Rattus norvegicus*) used for primary neuron collection were housed singly in chambers equipped with environmental enrichment. All studies were designed to minimize animal use and suffering. Rats were cared for by the UM Unit for Laboratory Animal Medicine. All individuals caring for animals were trained and approved in the care and long-term maintenance of rodent colonies, in accordance with the NIH-supported Guide for the Care and Use of Laboratory Animals. All personnel handling the rats and administering euthanasia were properly trained in accordance with the UM Policy for Education and Training of Animal Care and Use Personnel. Euthanasia was fully consistent with the recommendations of the Guidelines on Euthanasia of the American Veterinary Medical Association.

### Culturing and transducing iPSC-derived Neurons and Motor neurons

iNeuron and iMotor neuron neural progenitors (NPs) were made from iPSCs using the cell lines described in **Table 1** as follows. On DIV0, iPSCs were washed in PBS and incubated in accutase (Sigma, A6964) at 37°C for 5m. One volume of TeSR-E8 (StemCell, 059910) was added to the plate, and the cells were harvested and centrifuged at 300xg for 5m. The supernatant was removed and the pellet was resuspended in 1mL of fresh TeSR-E8 media containing ROCK inhibitor (Cayman Chemical, 10005583). Cells were then counted and 1.5X10^6^ cells were plated in TeSR-E8 media with ROCK inhibitor on a 60mm vitronectin (Fisher, A14700)-coated plate and incubated at 37°C overnight (O/N). On DIV1 and DIV2, media was replaced with TeSR-E8 containing 2µg/mL doxycycline. On DIV3, cells were harvested with accutase as above, frozen in TeSR-E8 with 10% DMSO at the desired cell number and stored in liquid nitrogen.

**Table 1.** Cell lines.

| **Name** | **Source** | **Additional information** | **Application** |
| --- | --- | --- | --- |
| HEK293T | ATCC | Identifier: CRL-3216;<br>RRID:CVCL_0063 | Roadblock-qPCR (**Figure 1f**), OPL Western blots (**Supplemental Figure 3b-c, e-f**), HaloLink and HaloTrap (**Supplemental Figure 4**), ICC (**Supplemental Figure 5**), DSG-crosslinking (**Supplemental Figure 8a-b**), CFTR assay (**Supplemental Figure 8c-f**) |
| Flp-In GFP-TDP43 HeLa cells | Gift from Clotilde Lagier-Tourenne, Massachusetts General Hospital | Ling et al. 2010<sup>41</sup> | qRT-PCR (**Figure 1b**, **Supplemental Figure 1a**), ICC (**Figure 2d-e**) |
| sTDP43-mNG HEK293T | This paper | Created as described in the Supplemental Materials and Methods | ICC (**Figure 3d**, **Supplemental Figure 1e**, **Supplemental Figure 2**), Live cell imaging (**Figure 3e-f**) |
| Rodent primary neurons | University of Michigan Unit for Laboratory Animal Medicine | Derived as described in the Supplemental Materials and Methods | Splicing reporter (**Figure 3b**), OPL (**Figure 4b-e**, **Supplemental Figure 3a, d**), ICC (**Figure 5c**, **Supplemental Figure 6b**), Survival analysis (**Figure 5d-e**, **Supplemental Figure 6c-d**, **Supplemental Figure 7**) |
| M17 human neuroblastoma cells | ATCC | Identifier: CRL-2267, lot 70044801 | *STMN2* reporter live cell imaging (**Figure 6b-c**) |
| 1021 + hNgn1/2 (iNeuron) | Weskamp et al. 2020<sup>19</sup> | Differentiated as described in the Supplemental Materials and Methods | qRT-PCR (**Figure 1c**), ICC (**Figure 2f-g**), Survival (**Figure 6e-f**) |
| 1021 + hNIL (iMotor neuron) | Chua et al. 2022<sup>48</sup> | Differentiated as described in the Supplemental Materials and Methods | qRT-PCR (**Figure 1d**) |
| WTC11 wild type iNeurons | Michael Ward Lab, NIH | Tian et al. 2019<sup>113</sup><br>Brown et al. 2022<sup>11</sup><br>Snyder et al. 2023<sup>106</sup><br>Seddighi et al. 2024<sup>114</sup> | HCR FISH (**Figure 6g-i**) |
| TDP43 CRISPRi iNeurons | Michael Ward Lab, NIH | Tian et al. 2019<sup>113</sup><br>Brown et al. 2022<sup>11</sup><br>Snyder et al. 2023<sup>106</sup><br>Seddighi et al. 2024<sup>114</sup> | HCR FISH (**Figure 6g-i**) |

iNeuron NPs were thawed and differentiation was performed as described previously^48–49^. Specifically, iNeuron NPs were thawed in TeSR-E8 media containing 2µg/mL doxycycline and ROCK inhibitor on Poly-L-ornithine (Sigma, P3655) and laminin (Sigma, L2020)-coated plates. On DIV1, media was changed to N2 media (TeSR-E8 media containing 1x N2 Supplement (Gibco, 17502-048), 1x NEAA Supplement (Gibco 11140-050), 10ng/mL BDNF (Peprotech, 450-02), 10ng/mL NT3 (Peptrotech 450-03), 0.2µg/mL laminin (Sigma L2020), 2mg/mL doxycycline). On DIV2, media was changed to a transition media (1x N2, 1x NEAA, 10ng/mL BDNF, 10ng/mL NT3, 0.2µg/mL laminin, 2mg/mL doxycycline in half TeSR-E8 media, half DMEM/F12 (Gibco, 11320-033)). On DIV3, media was changed into B27 media (1x B27, 1x Glutamax, 10ng/mL BDNF, 10ng/mL NT3, 0.2µg/mL laminin, and 1x Culture One (Gibco A33202-01) in Neurobasal-A media (Gibco, 12349-015). iNeurons were transduced on DIV3 with concentrated virus (prepared by the University of Michigan Vector Core) and sustained in the same culture medium for the remainder of the experiment. For qRT-PCR (**Figure 1**) and ICC experiments (**Figure 2**), cells were transduced for 7d, and 11j (SMG1i) and DMSO were added 48h prior to cell fixation/collection. For survival experiments (**Figure 6**), doxycycline was omitted from the DIV3 B27 media and imaging began on DIV6 and proceeded for 10d.

iMotor neuron NPs were thawed in TeSR-E8 containing 2µg/mL doxycycline and ROCK inhibitor on Matrigel (Sigma, CLS356234) or Poly-L-ornithine and laminin-coated plates. On DIV1, the media was changed to TeSR-E8 containing 2µg/mL doxycycline, 1x N2 supplement, 1:10,000 Compound E (Sigma, 565790). On DIV3, the media was changed to DMEM F12, 1X N2 supplement, 1x NEAA, 1x Glutamax, 2µg/mL doxycycline, and 1:10,000 Compound E. On DIV6, the media was changed to Neurobasal-A, 1x B27, 1x Glutamax, 10ng/mL BDNF, 10ng/mL NT3, 0.2 µg/mL mouse laminin (Sigma, L2020), 2µg/mL dox, and 1:100 Culture 1. For qRT-PCR experiments (**Figure 1**), iMotor neurons were transduced on DIV6 with concentrated virus (prepared by the University of Michigan Vector Core) for 6d, and 11j and DMSO were added 48h prior to cell collection.

### CRISPR/Cas9 integration of mNeonGreen into *sTARDBP* locus in HEK293T cells

sTDP43-mNG HEK293T cells were created using a dual-nickase CRISPR-Cas9 strategy^49,105^ as follows. First, oligos complimentary to the sequences flanking the region immediately upstream of the sTDP43 termination codon were annealed, digested, and inserted into the BbsI site of pX335, encoding Cas9n and chimeric single guide (sg)RNA (Addgene plasmid #188975; **Table 6**). HEK293T cells were then transfected with 1.25µg of each Cas9n-sgRNA pair, along with 2.5µg of a homology-directed repair cassette encoding the mNG open reading frame flanked by 400 bp of sequence homologous to the regions up and downstream of the target region. Following transfection, cells were split at a low density, allowing transfected cells to form individual colonies. Positive cells were identified by the presence of cytoplasmic green fluorescence and carefully transferred to a new dish. This process was repeated until 100% of cells displayed cytoplasmic mNG signal, and correct integration of the mNG cassette into *sTARDBP* locus was confirmed by PCR and Sanger sequencing.

### RT-PCR

RNA was isolated using the Direct-zol RNA Miniprep Kit (Zymo, R2052) and cDNA was reverse transcribed from 1µg of the resultant RNA with the High-Capacity cDNA Reverse Transcription Kit (Applied Biosystems, 4368814) in a reaction volume of 20µL. PCR for the CFTR assay (**Figure 6**) was performed using 1µL of cDNA as a template, GoTaq DNA Polymerase (Promega, M3001), and primers described in **Table 5**.

### qRT-PCR

RNA was isolated using the Direct-zol RNA Miniprep Kit or by phenol-chloroform extraction (described below). cDNA was reverse transcribed from 0.5-2µg of the resultant RNA with the High-Capacity cDNA Reverse Transcription Kit in a reaction volume of 20µL. 1µL of cDNA was used for each reaction as a template for quantitative (q)PCR, which was performed using Power SYBR Green (Applied Biosystems, A25742) using the primers listed in **Table 5**.

### Roadblock-qPCR

HEK293T cells were incubated with 400µM 4-thiouridine (4sU; Sigma, T4509) plus DMSO or 1µM 11j (SMG1i) for 0, 16 or 24h. After the incubation period, the cells were washed with ice cold PBS and collected in Trizol Reagent (Invitrogen, 15596018). RNA was isolated using the Direct-zol RNA Miniprep Kit. 1µg of resultant RNA was denatured at 65°C for 5m at a final volume of 11µL. Following denaturation, 10µL N-Ethylmaleimide (NEM) buffer (125mM Tris-HCl pH 8.0, 2.5mM EDTA, RNAse-free water), 3µL of 50 µg/µL NEM (Sigma, E3876), and 1µL of *in-vitro* transcribed firefly luciferase mRNA (0.2ng/µL) were added to each sample and incubated at 42°C for 1.5h. Next, 250ng of the NEM-modified RNA was reverse transcribed as follows: 1µL 50µM oligoDT was added to each sample and incubated at 60°C for 5m to anneal polyadenylated RNA. Then, 4µl 5X Protoscript II Buffer, 1µl Protoscript II Reverse Transcriptase (ProtoScript II Reverse Transcriptase, New England Biolabs, NC0319405), 1µL 10mM dNTPs (High-Capacity cDNA Reverse Transcription Kit, Applied Biosystems, 4368814), and 1µL 0.1M dithiothreitol (DTT; Thermo, R0862) were added to each sample and incubated at 42°C for 1h, heat inactivated at 65°C for 20m, and held at 4°C. 1µL of cDNA was used for each reaction as a template for qPCR, which was performed as described above.

### RNA Immunoprecipitation with HaloTrap

HEK293T cells were transfected with Halo-tagged constructs of interest (**Table 6**) 24h prior to UV crosslinking. To UV crosslink, HEK293T media was replaced with 1X PBS and cells were irradiated with 245nm UV at 150mJ/cm2 on ice. Cells were then harvested and pelleted at 235xg at 4°C for 10m, followed by a 30m lysis on ice with RNA lysis buffer (50mM Tris-HCl p.H 7.5, 150mM NaCl, 1% Triton-X100, 0.1% sodium deoxycholate, 1X protease inhibitor cocktail, RNAse inhibitor, DNAse). Lysates were then cleared at 4°C at 17000xg for 10m, and supernatants transferred to new tubes. 25µL input was saved for qRT-PCR. HaloTrap agarose beads (Chromotek, ota-20) were equilibrated as follows: 25µL beads per sample were diluted in 500µL cold dilution buffer (10mM Tris-HCL pH 7.5, 150mM NaCl, 0.5mM EDTA) and spun down at 2500xg for 5m before discarding supernatant. For protein binding to beads, the remaining 75µL of lysate was added to the equilibrated beads plus 300µL dilution buffer and incubated at 4°C for 1h with spinning. After incubation, samples were centrifuged at 2500xg for 5m and supernatant was collected for analysis of protein binding. Samples were then washed three times in 500µL dilution buffer supplemented with 0.05% IGEPAL (Thermo, J61055-AE) and the last wash was saved for further analysis. Phenol-chloroform extraction was performed to elute RNA as follows: first, Trizol Reagent was added directly to the beads and incubated on ice for 5 minutes prior to adding ⅕ volume of Chloroform (Thermo Fisher, AM9720) and vortexed until homogenous. Samples were then centrifuged at 4°C at 17000xg for 20 minutes. The aqueous phase was then carefully removed and transferred to a new tube. To precipitate the RNA, 75µg/mL glycogen (Thermo Fisher, R0551), 3M sodium acetate (pH 5.5, Fisher, AM7490) and 500µL of isopropanol (Fisher, A451-4) were added to the samples and incubated at -20°C O/N. The next day, samples were pelleted at 4°C at 17000xg for 20 minutes and washed twice with 75% EtOH and twice with 100% EtOH before drying completely in a vacufuge (Eppendorf Concentrator, 5301). The dried pellets were then DNAse treated (Baseline-ZERO DNase, Lucigen, DB0715K) for 30 minutes at room temperature (RT) and purified using the Monarch RNA Cleanup Kit (New England Biolabs, T2040). qRT-PCR was performed as described above.

### HCR FISH

HCR FISH custom probes were designed using Molecular Instruments Custom Probe Design Tool to target native and cryptic exons in *STMN2*^106^ (**Table 7**). HCR FISH was performed according to the manufacturer’s protocol^107^. Briefly, wildtype, sTDP43-transduced, and CRISPRi TDP43 knockdown iPSC-derived neurons (**Table 1**) were differentiated according to a previously published protocol^51^. On DIV3 after doxycycline, cells were re-plated on a 384-well plate (Corning) at a density of 3,000 cells per well in 100µL media. On DIV7 after doxycycline, cells were fixed with 4% paraformaldehyde for 10m at RT, then washed three times with PBS. Fixed cells were permeabilized with 70% ethanol O/N at -20°C. Ethanol was removed the following day and cells were washed twice with 2x SSC buffer (Molecular Instruments). Before adding the probe solutions, cells were incubated in warm probe hybridization buffer (Molecular Instruments) for 30m at 37°C. Probe solutions were prepared with 0.8 pmol of each probe set in probe hybridization buffer. Cells in probe solutions were hybridized O/N at 37°C. The following day, cells were washed four times with warm probe wash buffer. Cells were then washed twice with 5x SSCT (Molecular Instruments) at RT. Samples were amplified for 30m at RT in amplification buffer (Molecular Instruments) while hairpin solutions were prepared. 12 pmol of each hairpin one and two were heated to 95°C for 90s then cooled to RT without exposure to light for 30m. Cooled hairpins were added together in amplification buffer at RT before being added to the cells. Samples were incubated at RT O/N without exposure to light. The next day, hairpin solutions were removed, and cells were washed five times with 5x SSCT. Cells were incubated with Hoechst (Invitrogen, H3569) in PBS for nuclear counterstaining at a 1:10,000 dilution for 5m at RT. Cells were washed three times with PBS and stored at 4°C until being imaged. Cells were imaged with a Nikon spinning disk confocal on a 60x water immersion objective lens using a random imaging job with the nucleus as the plane of focus. NIS elements general analysis 3 was used to quantify the number of puncta (corresponding to native or cryptic exons), normalized to the number of nuclei per well.

### Capillary electrophoresis

Capillary electrophoresis was performed using the DNA 1000 Kit for 2100 Bioanalyzer Systems (Agilent, 5067-1504) according to manufacturer’s instructions.

### Immunocytochemistry

Cells were fixed with 4% paraformaldehyde (PFA; Sigma, P6148) for 10m, rinsed with PBS, and permeabilized with 0.1% Triton X-100 (Bio-rad, 161-0407) for 20m. Cells were then blocked in 3% bovine serum albumin (BSA; Fisher, BP9703-100) in PBS at RT for 1h before incubation O/N at 4°C in primary antibody diluted in 3% BSA. Cells were then washed 3 times in PBS and incubated at RT with secondary antibodies diluted 1:250 in 3% BSA for 1h. Following 3 washes in PBS containing 1:10,000 Hoechst 33258 dye (Invitrogen, H3569), cells were mounted on Superfrost Plus Microscope Slides (Fisher, 1255015) with Prolong Gold Antifade Mounting Reagent (Fisher, P10144) and imaged as described below. Images were analyzed by collecting measurements within cellular regions of interest using custom ImageJ/FIJI macros. Graphs were made using GraphPad Prism (Prism 9 for macOS).

### Western blotting

HEK293T and N2a cells were collected in PBS and pelleted at 10,000xg for 5m at 4°C and lysed in RIPA Buffer (Fisher, 29900) + Protease Inhibitor Cocktail (cOmplete, Mini, EDTA-free Protease Inhibitor Cocktail, Millipore, 118361700021) on ice for 30m. Lysates were sonicated at 80% amplitude with 5s on/5s off for a total of 1m using a Fisher Brand Model 50 Sonic Dismembrator and then centrifuged at 21,000g for 15m at 4°C. Supernatants were transferred to fresh tubes and equal volumes of each sample across conditions was diluted in 10x sample buffer (10% SDS, 20% glycerol, 0.0025% bromophenol blue, 100mM EDTA, 1M DTT, 20mM Tris, pH 8.0) and boiled for 8m. Samples were loaded onto a 10% SDS-PAGE gel with stacking gel and run at 110V. The blots were then transferred at 100V at 4°C for 1h onto an activated Immobilon-FL PVDF Membrane (Millipore, IPFL0010), blocked with 3% BSA in 0.2% Tween-20 (Sigma P9614) in Tris-buffered saline (TBST) for 20m, and blotted O/N at 4°C with primary antibody (**Table 4**) in 3% BSA in TBST. The following day, blots were washed 3 times in TBST, incubated at RT for 1h in secondary antibodies (**Table 4**) diluted 1:8000 in 3% BSA in TBST. Blots were then washed with TBST 3 times and imaged using an Odyssey CLx System (LI-COR). Image Studio was used to quantify bands.

### Immunoprecipitation with HaloLink

Immunoprecipitation using HaloLink was performed as described previously^19^. In summary, HEK293T cells were transfected with Halo-tagged constructs of interest (**Table 6**). After 48h, cells were collected in PBS and pelleted at 21,000xg. Each pellet was lysed in 100µL lysis buffer (50mM Tris-HCl, 150 mM NaCl, 1% Triton X-100, 0.1% sodium deoxycholate) on ice for 15m. 10µL of lysate input was saved for each sample. The remaining 90µL of lysate was added to 100µL of pre-washed HaloLink resin (Promega, 1914) which was prepared by washing and centrifuging for 2m at 800xg 3 times in wash buffer (100mM Tris pH 7.5, 150mM NaCl, 1mg/ml BSA, 0.005% IGEPAL). Each sample received an additional 400uL wash buffer prior to protein binding which took place on a tube rotator for 30m at RT. Samples were centrifuged at 800xg for 2m, and the supernatant was saved for further analysis. The resin was then washed 3 times in wash buffer and resuspended in elution buffer (1% SDS, 50 mM Tris-HCl Ph 7.5) and 10x sample buffer. Each sample was then boiled at 95°C for 10m and loaded onto a 10% SDS-PAGE gel alongside a fixed volume of input to assess protein binding. Western blot analysis was carried out as described above.

### DSG-crosslinking

HEK293T cells were transfected with EGFP-tagged constructs of interest 24h prior to DSG-crosslinking. Cells were collected, pelleted, and washed twice with 1x PBS and then resuspended in 100µL PBS with protease inhibitor cocktail. DSG or DMSO were added at a final concentration of 1mM and incubated at RT for 30m with shaking. Tris-HCl (pH 6.8) was added to each sample at a final concentration of 200mM to quench the DSG for 15m. Samples were centrifuged at 10,000xg for 5 min to obtain a pellet. Western blot analysis was carried out as described above.

### Confocal microscopy

Confocal images were taken on a Nikon AXR NSPARC confocal system with a 60x NA1.42 Oil/DIS PLan-Apochromat Lambda D objective with a working distance of 1.5mm, and a 40x CFI Apochromat LWD Lambda S objective with a working distance of 0.30mm.

### Longitudinal microscopy

Neurons were imaged as described previously^19,49,63–64^ using a Nikon Eclipse Ti inverted microscope with PerfectFocus3a 20X objective lens and either an Andor iXon3 897 EMCCD camera or Andor Zyla 4.2 (+) sCMOS camera. A Lambda CL Xenon lamp (Sutter) with 5mm liquid light guide (Sutter) was used to illuminate samples, and custom scripts written in Beanshell for use in micromanager controlled all stage movements, shutters, and filters. For automated analyses of primary neuron survival, custom ImageJ/FIJI macros and Python scripts were used to identify neurons and draw cellular regions of interest (ROIs) based upon size, morphology, and fluorescence intensity. Custom Python scripts were used to track ROIs over time, and cell death marked a set of criteria that include rounding of the soma, loss of fluorescence and degeneration of neuritic processes^64,108^. The same ROIs were used to track fluorescence intensity over time, contributing to the sTDP43 splicing reporter (**Figure 3**), OPL (**Figure 4**), and STMN2 reporter (**Figure 6**) findings. For manual analyses of iNeuron survival, image time-series were processed by flat-field correction and image registration followed by programmatic de-identification for blinded analysis. Time-series were uploaded to a browser-based server or CVAT^109^ where a trained user manually counted survival using the point tracking mode. Cell death was identified by deformation of rounded somas into amorphous blebs and was recorded by marking tracks as ‘outside’ at time of death. Tracks not marked as “outside” were right-censored. Lifelines^110^ was used to produce Cox proportional hazard baseline plots from the resulting data.

### Drosophila studies

Homozygous male flies containing the EGFP-tagged transgene of interest were crossed to gmr-Gal4 (**Table 2**) females at 29°C on SY10 food. Progeny eyes were imaged at day 1-3 post eclosion. Rough eye phenotype was assessed by an experimenter blinded to genotype using previously published methods^111–112^. Fly images were taken using a Leica M125 stereomicroscope and a Leica K3C equipped with LAS X manual z-stacking software. A minimum of 30 flies were analyzed per cross and data was derived from three independent experiments.

**Table 2.** *Drosophila* lines.

| Name | Source | Identifiers and Additional Information |
| --- | --- | --- |
| uAS-EGFP | Peter Todd<br>lab, University<br>of Michigan | BDSC #5431 |
| uAS-EGFP-sTDP43 | This paper | BestGene *Drosophila* Embryo Injection<br>Services<br>PhiC31 source: P{nos-phiC31\|int.NLS}X<br>Insertion site: ZH-86Fb |
| uAS-EGFP-TDP43 | This paper | BestGene *Drosophila* Embryo Injection<br>Services<br>PhiC31 source: P{nos-phiC31\|int.NLS}X<br>Insertion site: ZH-86Fb |
| gmr-Gal4 | Peter Todd<br>Lab, University<br>of Michigan | BDSC #8605 |
| tub5-GS | Scott Pletcher<br>Lab, University<br>of Michigan | Osterwalder et al. 2001<sup>115</sup> |
| w1118 | Peter Todd<br>Lab, University<br>of Michigan | BDSC #3605 |

**Table 3.**
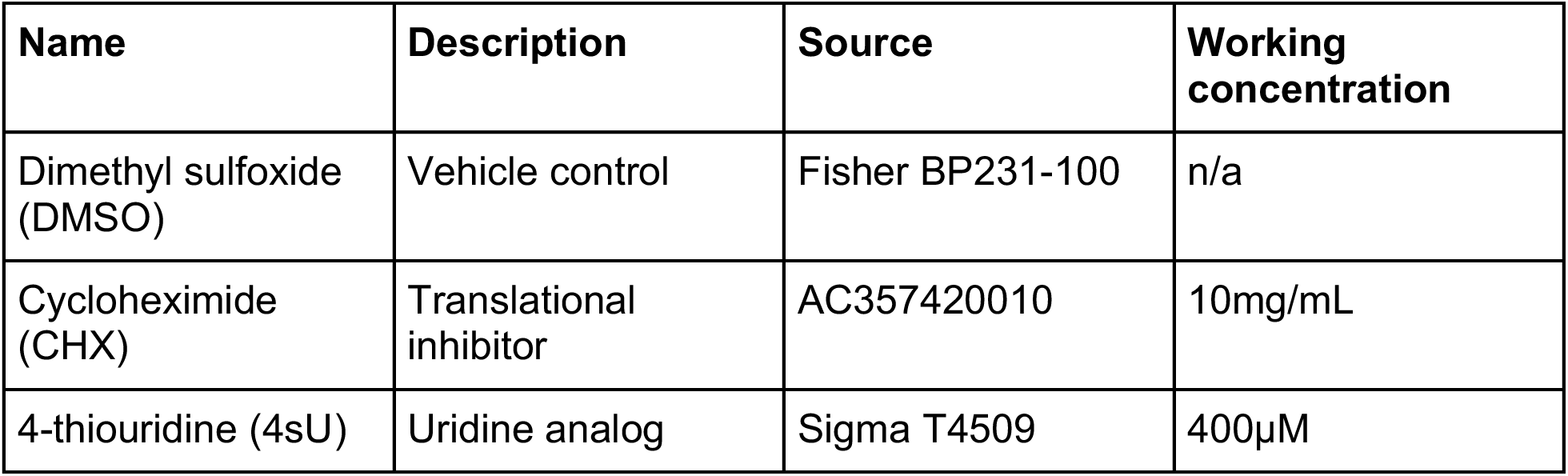
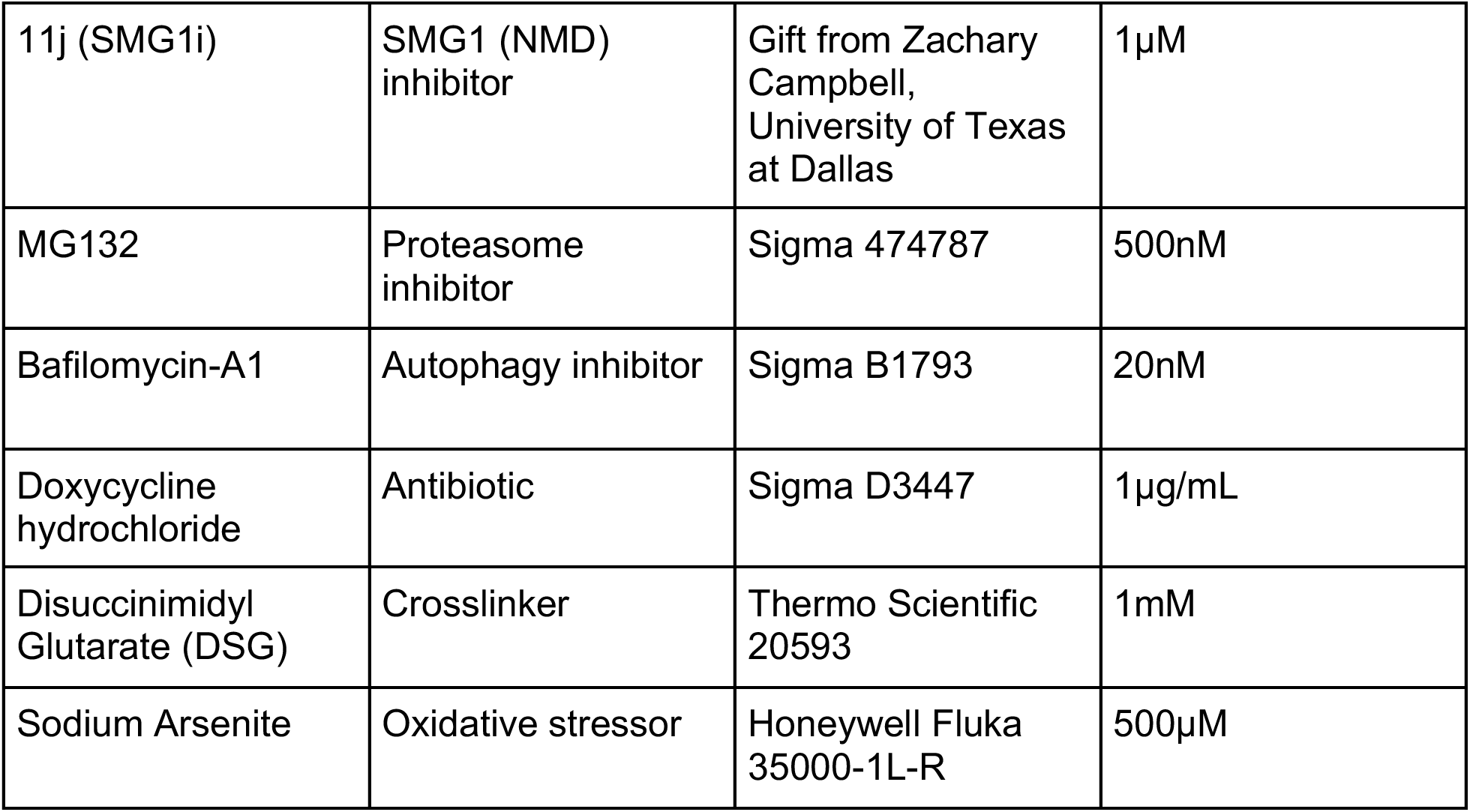
Drugs.

| **Name** | **Description** | **Source** | **Working**<br>**concentration** |
| --- | --- | --- | --- |
| Dimethyl sulfoxide<br>(DMSO) | Vehicle control | Fisher BP231-100 | n/a |
| Cycloheximide<br>(CHX) | Translational<br>inhibitor | AC357420010 | 10mg/mL |
| 4-thiouridine (4sU) | Uridine analog | Sigma T4509 | 400µM |
| 11j (SMG1i) | SMG1 (NMD) inhibitor | Gift from Zachary Campbell, University of Texas at Dallas | 1μM |
| MG132 | Proteasome inhibitor | Sigma 474787 | 500nM |
| Bafilomycin-A1 | Autophagy inhibitor | Sigma B1793 | 20nM |
| Doxycycline hydrochloride | Antibiotic | Sigma D3447 | 1μg/mL |
| Disuccinimidyl Glutarate (DSG) | Crosslinker | Thermo Scientific 20593 | 1mM |
| Sodium Arsenite | Oxidative stressor | Honeywell Fluka 35000-1L-R | 500μM |

**Table 4.** Antibodies.

| **Antibody** | **Source** | **Catalog number** | **Species** | **Dilution** |
| --- | --- | --- | --- | --- |
| ***Primary*** |  |  |  |  |
| sTDP43 (5D32) | Custom-made from Genscript and synthesized by Peter Tessier's Lab, University of Michigan | NA | Mouse | 1:250 for ICC |
| sTDP43 (734) | Weskamp et al. 2020<sup>19</sup> | NA | Rabbit | 1:250 for ICC<br>1:100 for WB |
| sTDP43 (7689) | Custom-made from Genscript | NA | Rabbit | 1:250 for ICC |
| MAP2 | Novus | NB300-213 | Chicken | 1:1000 for ICC |
| pUb | Cell Signaling Technology | 58395 | Rabbit | 1:2000 for WB |
| P62 (SQSTM1) | Abnova | H00008878-M01 | Mouse | 1:1000 for WB |
| GAPDH | Millipore Sigma | MAB374 | Mouse | 1:1000 for WB |
| GFP (mAb) | Invitrogen | MA5-15256 | Mouse | 1:1000 for WB |
| β-Actin | Cell Signaling Technology | 4967S | Rabbit | 1:1000 for WB |
| mNeonGreen 32F6 | Chromotek | 32f6-100 | Mouse | 1:500 for ICC |
| TDP43 (C-terminal) | Proteintech | 12892 | Rabbit | 1:500 for ICC |
| TDP43 (RRMs) | Proteintech | 10782 | Rabbit | 1:1000 for WB |
| GFP (pAb) | Abcam | ab290 | Rabbit | 1:1000 for WB |
| **Secondary** |  |  |  |  |
| Alexa Fluor Plus 488 | Invitrogen | A32790TR | Donkey anti rabbit | 1:250 |
| Alexa Fluor 546 | Invitrogen | A11035 | Goat anti rabbit | 1:250 |
| Alexa Fluor 546 | Invitrogen | A11030 | Goat anti mouse | 1:250 |
| Alexa Fluor 647 | Invitrogen | A31571 | Donkey anti mouse | 1:250 |
| Alexa Fluor 647 | Invitrogen | A-21449 | Goat anti-chicken | 1:250 |
| Alexa Fluor 647 | Invitrogen | A31573 | Donkey anti rabbit | 1:250 |
| LICOR IRDye 800CW | Licor | 926-32210 | Goat anti mouse | 1:8000 |
| LICOR IRDye 680RD | Licor | 926-68073 | Donkey anti rabbit | 1:8000 |
| LICOR IRDye 800CW | Licor | 926-32213 | Donkey anti rabbit | 1:8000 |
| LICOR<br>IRDye 680RD | Licor | 926-68072 | Donkey anti<br>mouse | 1:8000 |

For immunoblotting, 10 males and 10 females from the F1 progeny from the cross described above were included in each sample. The heads of each fly were removed with a razor blade, lysed in 100µL RIPA buffer with protease inhibitor, hand homogenized (BioSpec Products, 1083MC), passed through a 28.5G needle 8 times and incubated on ice for 15m. Samples were centrifuged at 4°C for 5m at 12000xg to pellet debris. Supernatants were transferred to fresh tubes and equal volumes of each sample across conditions was diluted in 10x sample buffer and boiled for 8m. Western blot was carried out as described above.

For qRT-PCR analysis of transgene expression, homozygous male flies containing the transgene of interest were crossed to female tub5-GS (**Table 2**) virgins at 25°C on SY10 food. The F1 progeny were placed on fresh SY10 + RU486 food for 72h to induce full-body transgene expression. 8 males and 8 females were included in each sample and RNA analysis was performed in triplicate. Each sample was homogenized in 500µL Trizol. RNA was isolated with phenol-chloroform extraction and qRT-PCR analysis was carried out as described above.

### Plasmids

Primers used for site-directed mutagenesis (SDM) and PCR amplifications were ordered from Integrated DNA Technologies (IDT) (**Table 5**). GeneBlocks were obtained from IDT and BlueHeron (**Table 6**). SDM was conducted using the Q5 Site-Directed Mutagenesis Kit (New England Biolabs, E0554) and PCR amplifications were carried out using PrimeSTAR GXL DNA Polymerase (Takara, R050A) according to manufacturer’s instructions. All synthesized plasmids were verified by Sanger sequencing and described in **Table 6**.

**Table 5.**
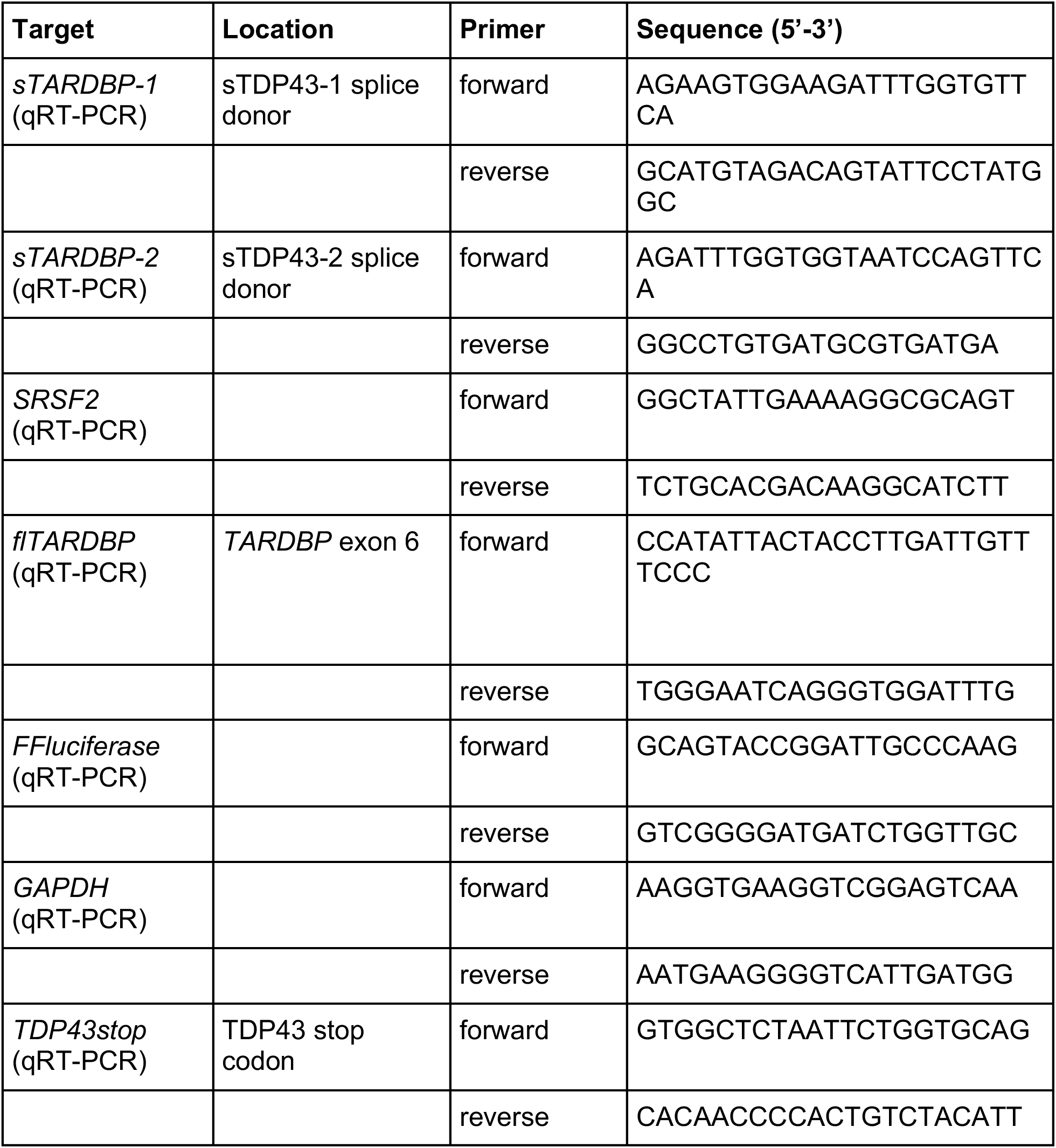
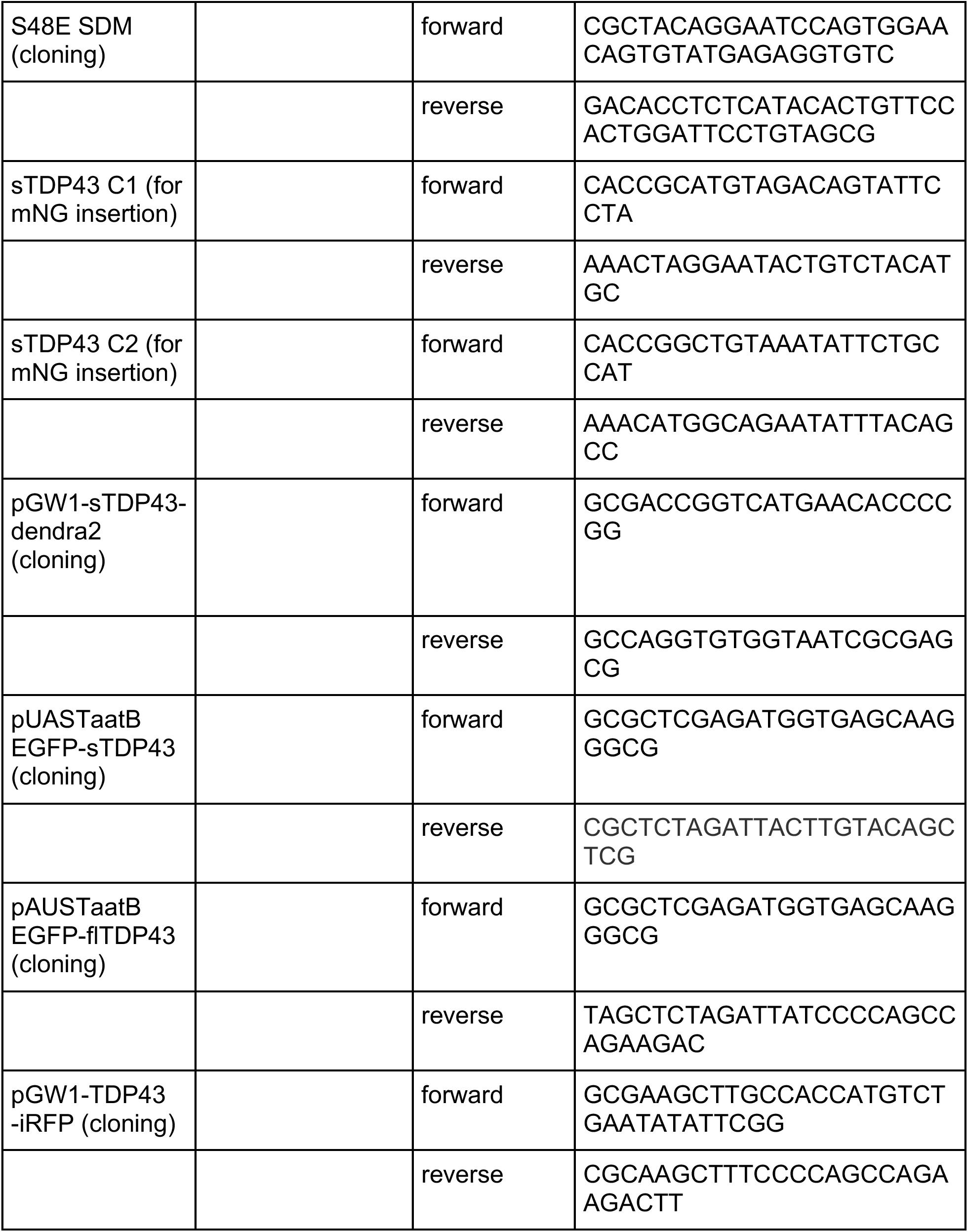
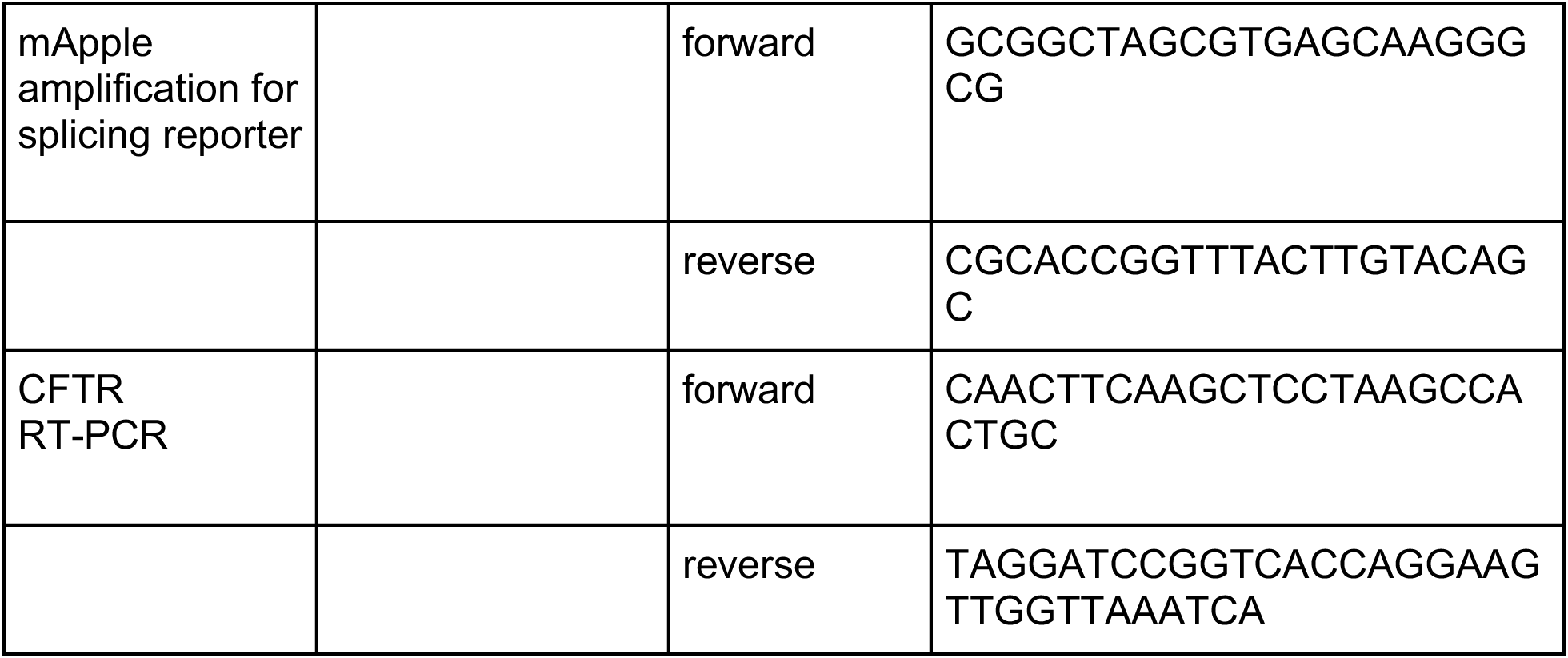
Primers.

**Table 6.**
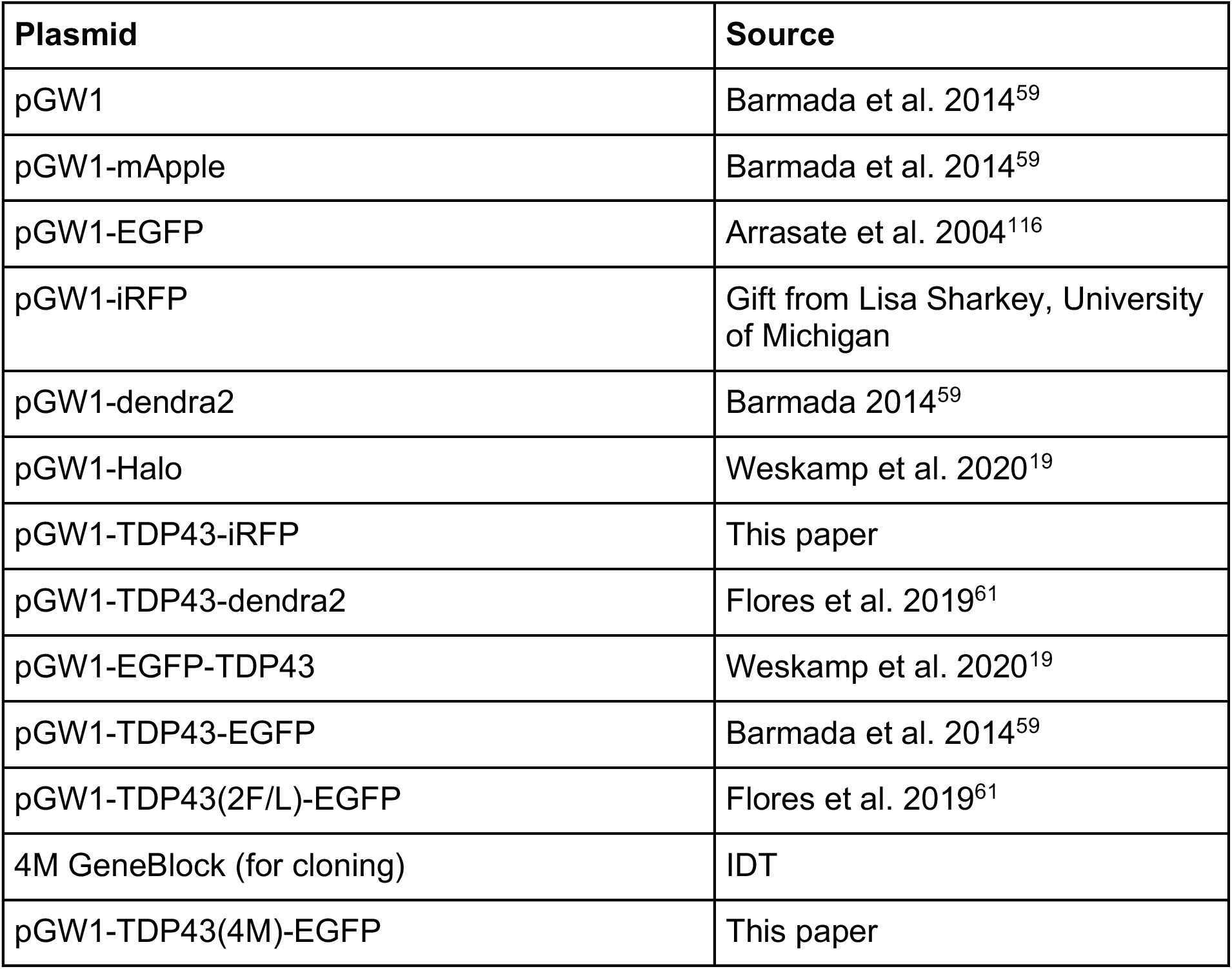
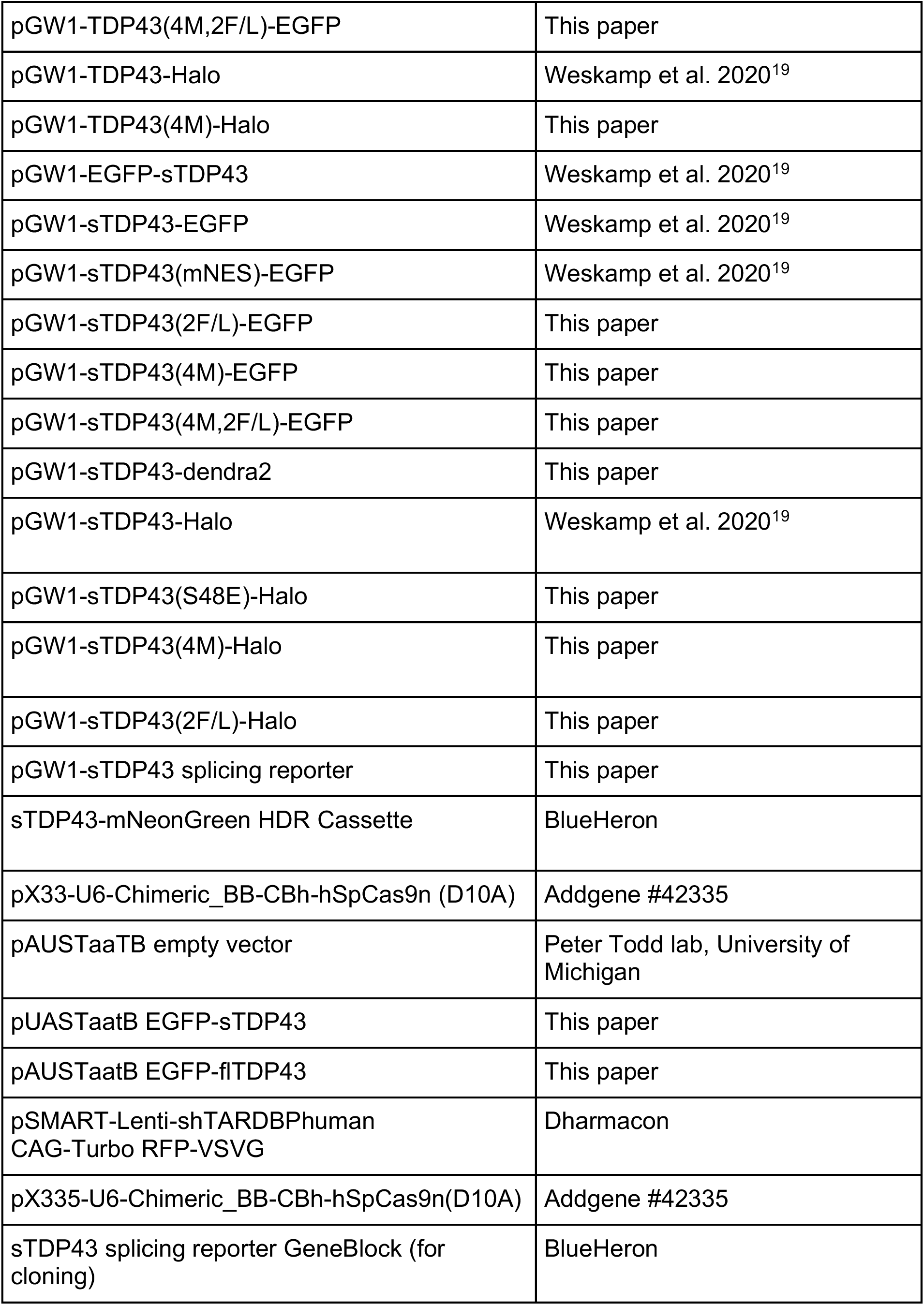
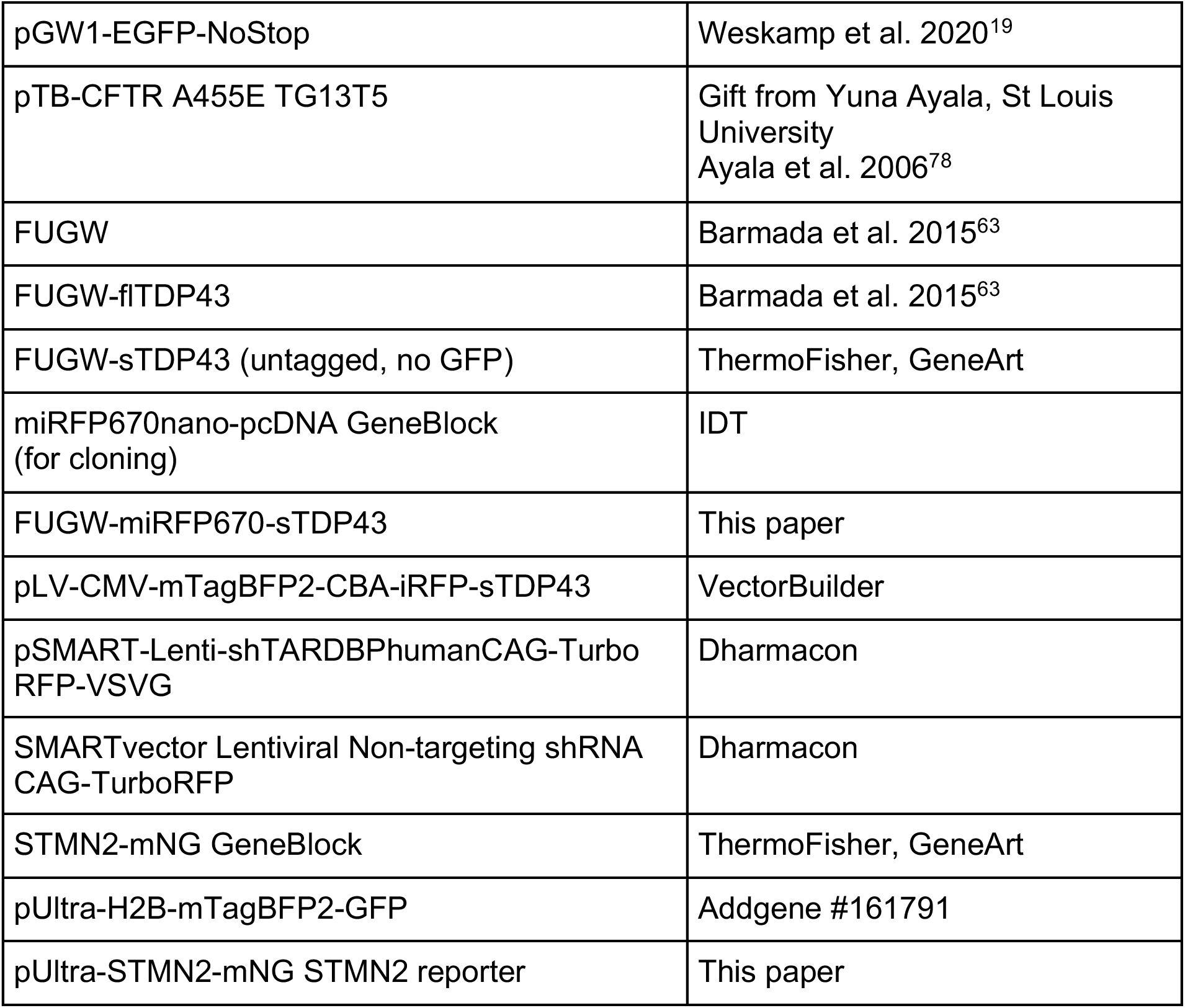
Plasmids.

**Table 7.**
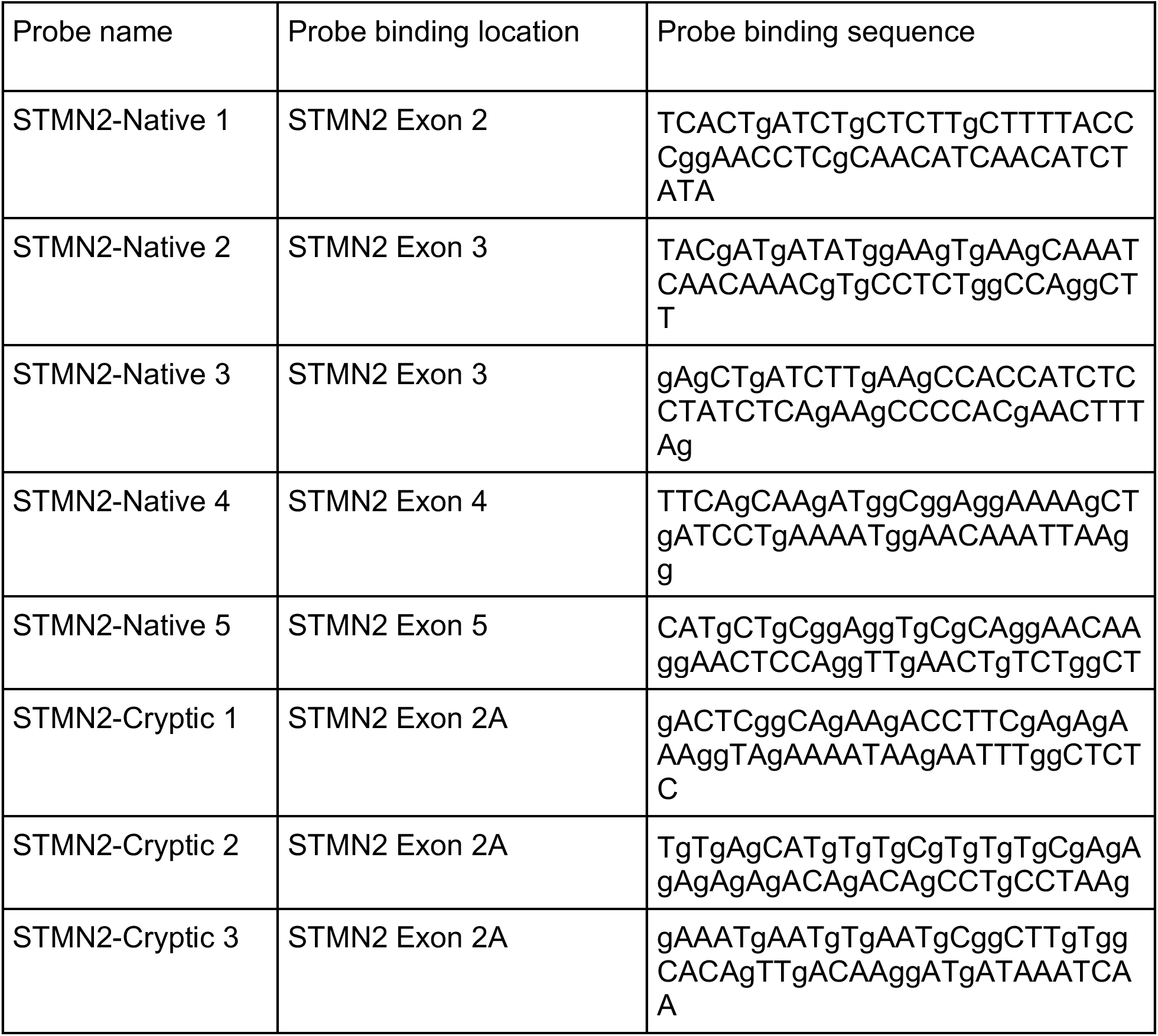
HCR-FISH probes.

To synthesize **pGW1-TDP43-iRFP**, pGW1-TDP43-EGFP was PCR amplified using primers (**Table 5**) adding HindIII sites flanking the TDP43 ORF. The resulting PCR product was gel isolated, digested with HindIII, and inserted into the cut and phosphorylated HindIII sites in pGW1-iRFP.

**pGW1-sTDP43(2F/L)-EGFP** was created using pGW1-sTDP43-EGFP as a template and the primers noted in **Table 5**.

To create **pGW1-sTDP43(4M)-EGFP** and **pGW1-TDP43(4M)-EGFP**, a GeneBlock comprised of the N-terminus of the TDP43 open reading frame containing 4M mutations (E17A, E21A, R52A, R55A) and flanked by BsteII and PpuMI restriction enzyme sites was generated by IDT. This GeneBlock was digested with BstEII and PpuMI and cloned into the corresponding sites in both pGW1-sTDP43-EGFP and pGW1-TDP43-EGFP vectors.

To make **pGW1-sTDP43(4M,2F/L)-EGFP** and **pGW1-TDP43(4M,2F/L)-EGFP**, pGW1-sTDP43(4M)-EGFP and pGW1-TDP43(4M)-EGFP were digested with PpuM1 and HindIII restriction enzymes to isolate small fragments of the N-terminus containing the 4M mutations. These fragments were then ligated into the corresponding sites in both pGW1-sTDP43(FL)-EGFP and pGW1-TDP43(FL)-EGFP.

**pGW1-sTDP43(2F/L)-Halo** was made by replacing the EGFP in pGW1-sTDP43(FL)-EGFP with HaloTag from pGW1-sTDP43-HaloTag using AgeI and SbfI.

To make **pGW1-sTDP43(4M)-Halo** and **pGW1-TDP43(4M)-Halo**, the N-terminal region containing the 4M mutations was cut from pGW1-sTDP43(4M)-EGFP using Kpn1 and PpUM1 and ligated into the corresponding sites of pGW1-sTDP43-HaloTag and pGW1-TDP43-HaloTag, respectively.

To make **pGW1-sTDP43-dendra2**, pGW1-dendra2 was PCR amplified using primers (**Table 5**) to add an AgeI site upstream and a NruI site downstream of the fluorescent tag. The PCR products were gel isolated, digested with AgeI and NruI, and ligated into pGW1-sTDP43-EGFP.

To create **pGW1-sTDP43(S48E)-Halo**, the S48E mutation was first introduced to pGW1-sTDP43-EGFP via SDM. Once this plasmid was sequence verified, it was digested with AgeI and SbfI to cut out the EGFP fusion at the C-terminus of sTDP43 and replaced with a HaloTag from pGW1-sTDP43-Halo.

To generate the **sTDP43 splicing reporter**, the mApple open reading frame was amplified from pGW1-mApple, adding an NheI site at the 5’ end and an AgeI site at the 3’ end. The resultant amplicon was gel isolated, digested with NheI and AgeI, and cloned into the corresponding sites of pGW1-EGFP-NoStop to create pGW1-EGFP-mApple. Next, a GeneBlock containing sTDP43 exon 6 and 3’UTR (BlueHeron) was digested with Kpn1 and NheI, then inserted into the same sites on pGW1-EGFP-mApple between the two fluorescent tags.

**pUASTaatB-EGFP-sTDP43** and **pUASTaatB-EGFP-flTDP43** were created first by PCR amplification of pGW1-EGFP-sTDP43 and pGW1-EGFP-TDP43, respectively, using the primers listed in **Table 5**. The pAUSTaaTB empty vector was then digested with XbaI and XhoI, and gel purified. The PCR products were digested with XbaI and XhoI before ligating into the cut pAUSTaaTB vector.

The ***STMN2* minigene reporter** (pUltra-STMN2-mNG) was created using a STMN2-mNG GeneBlock. This construct was digested with PvuI, EcoRI and AgeI, and cloned into the corresponding EcoRI and AgeI sites in a pUltra-H2B-mTagBFP2-GFP backbone in place of the mTagBFP2 and GFP.

**sTDP43-FUGW** was ordered from ThermoFisher GeneArt Gene Synthesis.

**FUGW-miRFP670-sTDP43** was cloned using a GeneBlock containing the miRFP670nano-pcDNA (Addgene #127443) open reading frame flanked by AgeI and BsteII digestion sites. The GeneBlock was digested with AgeI and BsteII and inserted into the corresponding sites upstream of sTDP43 in FUGW-sTDP43.

**pLV-CMV-mTagBFP2-CBA-iRFP-sTDP43** was ordered from VectorBuilder.

## Author contributions

MMD and SJB designed the study. MMD generated plasmids, performed RT-PCR, qRT-PCR, capillary electrophoresis, western blots, immunocytochemistry, confocal and automated microscopy, and conducted all corresponding analyses. KW generated plasmids. NBG assisted with microscopy analyses and cloning. JW wrote the code for image acquisition and neuronal survival analysis. ET generated plasmids, maintained iPSCs and differentiated them into neurons. MRG assisted with *Drosophila* husbandry and rough eye analysis. AS performed HCR FISH. EP and MB provided intellectual contributions and assisted with neuronal survival studies. XL isolated and cultured primary neurons. JB and SS immunostained and imaged cells by confocal microscopy. JN analyzed survival and microscopy data. CW, CS and SW helped clone, analyze, and validate sTDP43 constructs. NG and JJM conducted initial experiments on sTDP43 levels. JJM and AVS helped develop methods and contributed to fluorescence microscopy. MMD and SJB assembled figures, wrote and edited the manuscript.

## Figure legends

**Supplemental figure 1. sTDP43-specific antibodies confirm regulation by NMD and colocalize with mNG signal in knock-in HEK293T cells.**

**(a)** As detected by qRT-PCR, sTDP43 transcripts are induced by GFP-TDP43 overexpression and NMD inhibition by Cycloheximide (CHX) in Flp-In GFP-TDP43 HeLa cells. (**b**) Diagram of the NMD pathway. Premature termination codon (PTC)-containing transcripts are targeted for degradation following UPF1 phosphorylation by SMG1. CHX blocks translation, while 11j (SMG1i) inhibits SMG1. (**c**) Western blot of N2a cells overexpressing GFP-TDP43 and/or treated with SMG1i and probed with a sTDP43-specific antibody. (**d**) Quantification of sTDP43 immunoreactivity in (**c**), demonstrating significant increase in sTDP43 protein abundance with TDP43 OE and SMG1i. (**a**, **d**) Data combined from 3 biological replicates. *p<0.05, **p<0.01, ****p<0.0001 by 2-way ANOVA with Tukey’s multiple comparisons test. (**e**) Representative images demonstrating overlapping signals between endogenous sTDP43-mNG fluorescence, immunoreactivity from an anti-mNG antibody, and two distinct antibodies raised against sTDP43. Scale bars: 25µm.

**Supplemental figure 2. sTDP43-mNG puncta overlap minimally with cytosolic organelles.**

Confocal microscopy images of intrinsic sTDP43-mNG signal in comparison with the stress granule marker G3BP1 with (**a**) and without (**b**) sodium arsenite (NaAsO_2_) to induce the formation of stress granules, the processing (P)-body marker Xrn1 with (**c**) and without (**d**) NaAsO_2_ to induce stress granule formation, as well as the lysosomal marker Lamp1 (**e**) and the transport granule marker Stau1 (**f**). Scale bars: 10µm; inset scale bars: 5µm.

**Supplemental figure 3. sTDP43 is cleared by the UPS and macroautophagy.**

**(a)** Half-life of dendra2 and sTDP43-dendra2 expressed in rodent primary neurons treated with 500nM MG132. N= number of neurons combined among 4 experiments. Dendra2+vehicle n=201, dendra2+MG132 n=61, sTDP43-d2+vehicle n=360, sTDP43-d2+MG132 n=312. (**b**) Immunoblot and (**c**) quantification of polyubiquitin in HEK293T cells treated for 24h with increasing doses of MG132. (**d**) Half-life of dendra2 and sTDP43-dendra2 in rodent primary neurons treated with 20nM Bafilomycin-A1 (BafA1). N= number of neurons combined among 3 experiments. Dendra2+vehicle n=255, dendra2+BafA1 n=48, sTDP43-d2+vehicle n=395, sTDP43-d2+BafA1 n=35. (**e**) Western blot and (**f**) quantification of p62, an autophagy substrate, in HEK293T cells treated for 24h with increasing doses of BafA1. (**a**, **d**) Data combined and stratified among 3 biological replicates. *p<0.5, ****p<0.0001 by 1-way ANOVA and Tukey’s multiple comparisons test. (**c**, **f**) Data combined from 3 biological replicates. *p<0.05, **p<0.01 by 1-way ANOVA with Dunnett’s post-hoc test. (**g**) qRT-PCR validation of similar transgene expression at the RNA level for EGFP-flTDP43 and EGFP-sTDP43 in *Drosophila*. (**h**) Immunoblot and (**i**) quantification confirming relatively equivalent expression of EGFP-flTDP43 and EGFP-sTDP43 transgenes in *Drosophila*. *Nonspecific band. *p<0.05, ***p<0.001 by 1-way ANOVA with Dunnett’s.

**Supplemental figure 4. Engineered mutations impact the ability of sTDP43 variants to oligomerize and bind RNA.**

**(a)** Schematic of HaloTrap and HaloLink verification of the consequences of mutations on sTDP43 oligomerization and RNA binding. (**b**) Immunoblot demonstrating the disruption of sTDP43 oligomerization with itself (sTDP43-Halo) and endogenous (endo.fl)TDP43 by the 4M, but not S48E, mutations. (**c**) RNA immunoprecipitation showing impaired RNA binding in sTDP43 variants carrying the 2F/L mutation. Data combined from 3 biological replicates. **p<0.01, ***p<0.001 by 1-way ANOVA with Tukey’s.

**Supplemental figure 5. Mutations affecting RNA binding and dimerization of sTDP43 alter its subcellular localization.**

**(a)** Confocal microscopy images showing the distribution of each sTDP43-EGFP mutant in HEK293T cells. (**b**) Nuclear/cytoplasmic ratio for each variant in HEK293T cells. N= number of cells analyzed among 3 biological replicates: sTDP43-EGFP n=101, sTDP43(mNES)-EGFP n=83, sTDP43(2F/L)-EGFP n=110, sTDP43(4M)-EGFP n=94, sTDP43(4M,2F/L)-EGFP n=101. ****p<0.0001 by 1-way ANOVA with Dunnett’s multiple comparisons test. Scale bars: 10µm.

**Supplemental figure 6. The RNA binding and dimerization domains are essential for flTDP43-mediated toxicity.**

**(a)** Schematic depicting the location of flTDP43 mutations. (**b**) Representative images of rat primary cortical neurons expressing EGFP and each EGFP-tagged flTDP43 construct. Scale bars: 5µm. (**c**) Cumulative hazard and (**d**) forest plot of hazard ratios of flTDP43 variants expressed in rat primary cortical neurons, determined by automated fluorescence microscopy. N=number of neurons, combined and stratified among 6 biological replicates: EGFP n=2047, flTDP43-EGFP n=2034, flTDP43(2F/L)-EGFP n=540, flTDP43(4M)-EGFP n=480, flTDP43(4M,2F/L)-EGFP n=1378. The negative control (EGFP) serves as a reference (Hazard ratio=1). ***p<0.001 by Cox proportional hazard analysis.

**Supplemental figure 7. Bootstrapping of survival data confirms the impact of RNA binding, oligomerization and localization of sTDP43- and flTDP43-mediated toxicity.**

(**a,c**) Cumulative hazard plots of (**a**) sTDP43 and (**c**) flTDP43 variants, bootstrapped to include 100 randomly selected neurons from each experiment. EGFP n=577, sTDP43- EGFP n=600, sTDP43(4M)-EGFP n=297, sTDP43(2F/L)-EGFP n=300, sTDP43(4M,2F/L)-EGFP n=300, sTDP43(mNES)-EGFP n=300, flTDP43-EGFP n=300, flTDP43(4M)-EGFP n=300, flTDP43(2F/L)-EGFP n=300, flTDP43(4M,2F/L)-EGFP n=300. (**b**,**d**) Forest plots depicting cumulative hazard ratios of 10 different bootstrapped datasets, corresponding to data shown in (**a**) and (**c**). *p<0.05, ***p<0.001 by Cox proportional hazard analysis.

**Supplemental figure 8. sTDP43 affects endogenous flTDP43 function via oligomerization and RNA binding.**

sTDP43 interferes with the function of endogenous flTDP43 via its N-terminal oligomerization and RNA binding properties.

**(a)** Immunoblot and (**b**) quantification of DSG-crosslinking of sTDP43-EGFP variants, showing decreased oligomerization among 4M-containing sTDP43 mutants (sTDP43(2F/L)-EGFP, sTDP43(4M,2F/L)-EGFP) and nuclear sTDP43 (sTDP43(mNES)-EGFP) relative to WT sTDP43-EGFP. *p<0.05 by 1-way ANOVA with Dunnett’s multiple comparisons test, combined among 5 biological replicates. Pink bracket represents monomeric sTDP43-EGFP variants, yellow bracket represents oligomeric sTDP43-EGFP variants. (**c**) Schematic of cystic fibrosis transmembrane conductance regulator (CFTR) assay. TDP43-mediated splicing of the reporter drives exon 9 exclusion under normal conditions. Compromised TDP43 splicing causes cryptic splicing and exon 9 retention. Arrows indicate primers used to amplify splice junctions. (**d**) Capillary electrophoresis of CFTR splice products following expression of flTDP43-EGFP and sTDP43-EGFP variants in HEK293T cells. (**e**-**f**) Quantification of spliced (**e**), and cryptic products (**f**) in (**d**). Statistical analysis was conducted on log-transformed values. *p<0.05, ***p<0.001, ****p<0.0001 by 1-way ANOVA with Tukey’s multiple comparisons test. LOF: endogenous TDP43 loss-of-function, GOF: endogenous TDP43 gain-of-function

