## Supplemental Figures for "TDP43 autoregulation gives rise to shortened isoforms that are tightly controlled by both transcriptional and post-translational mechanisms"

### Supplemental Figure 1

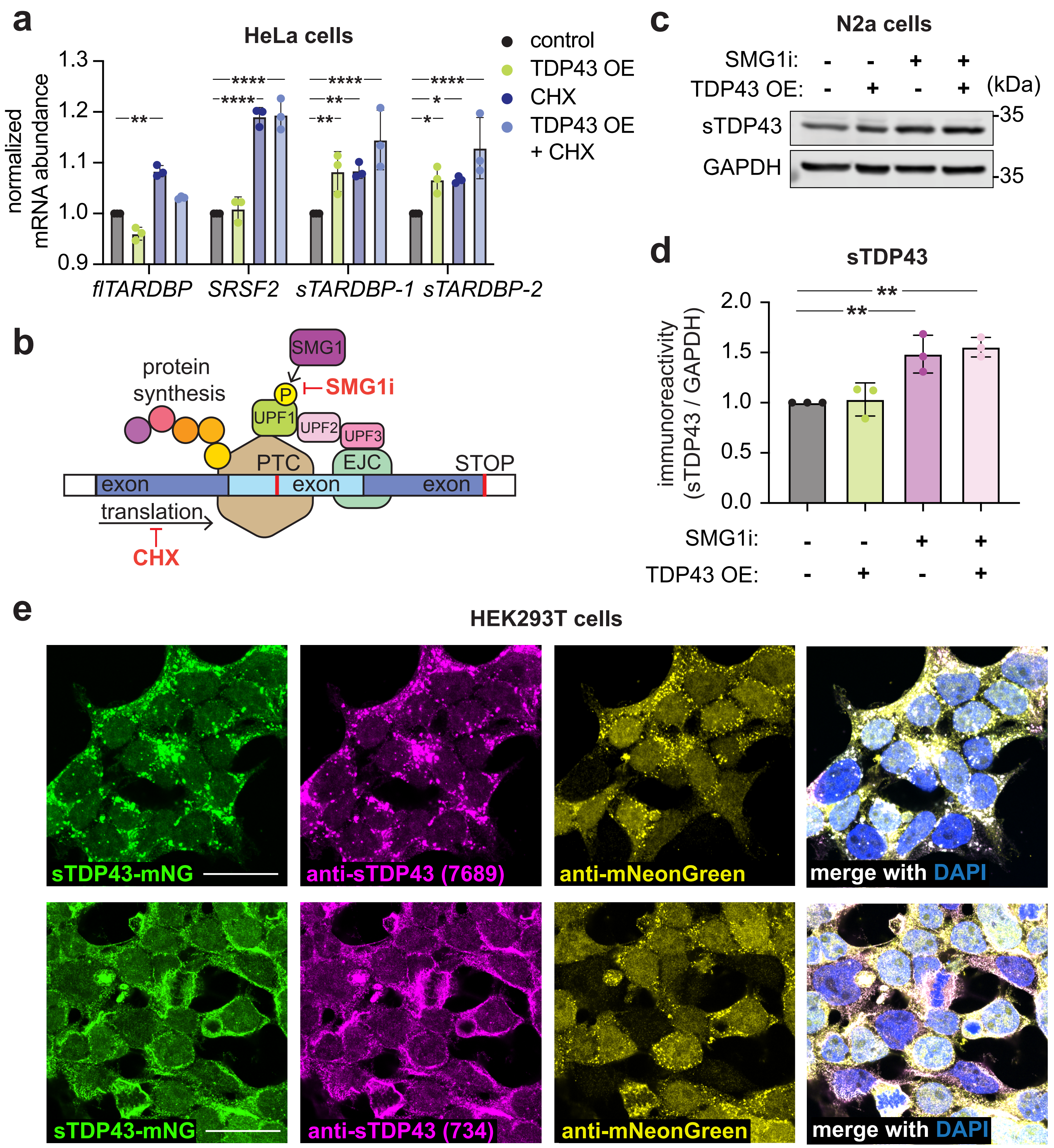

### Supplemental Figure 2

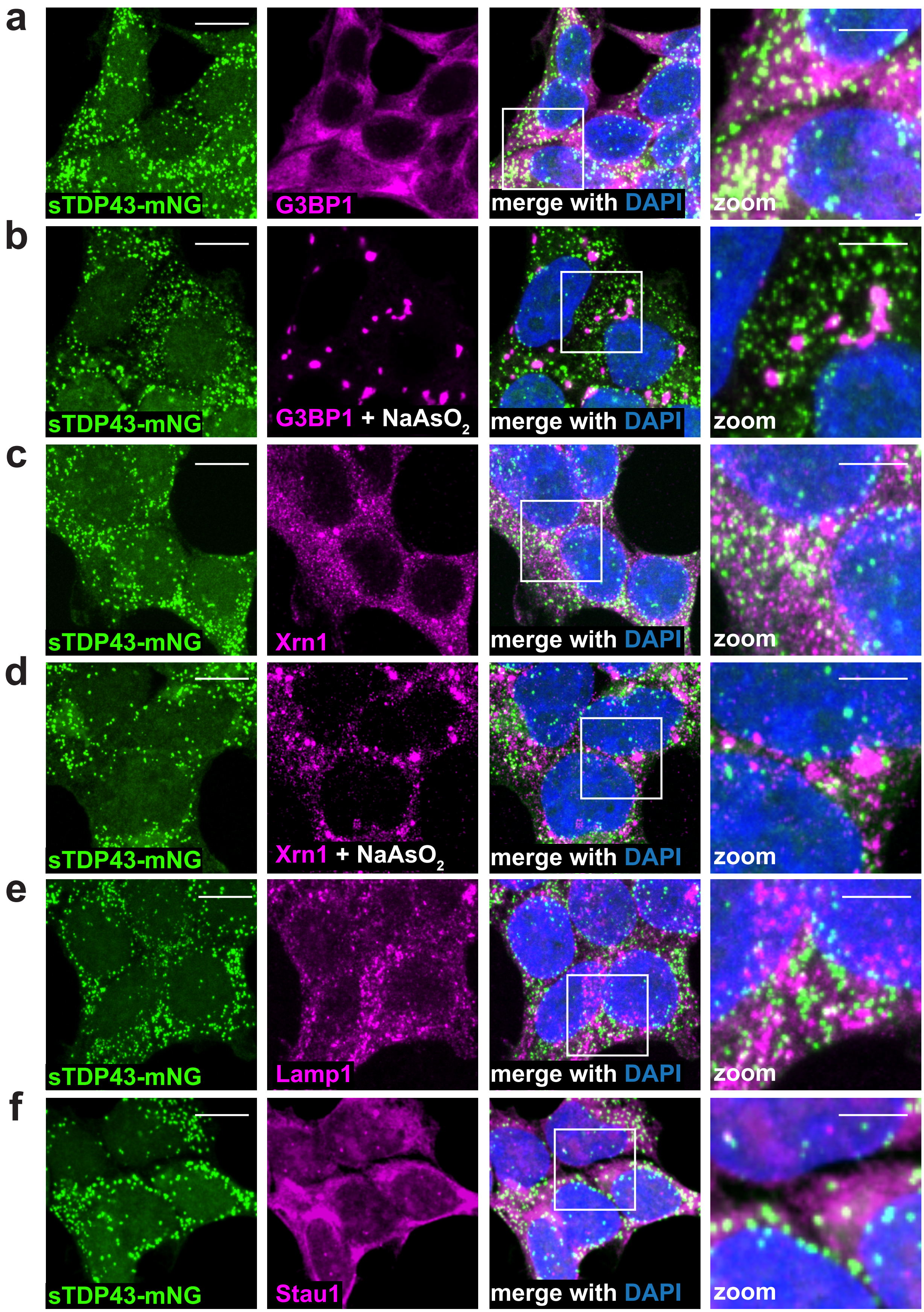

### Supplemental Figure 3

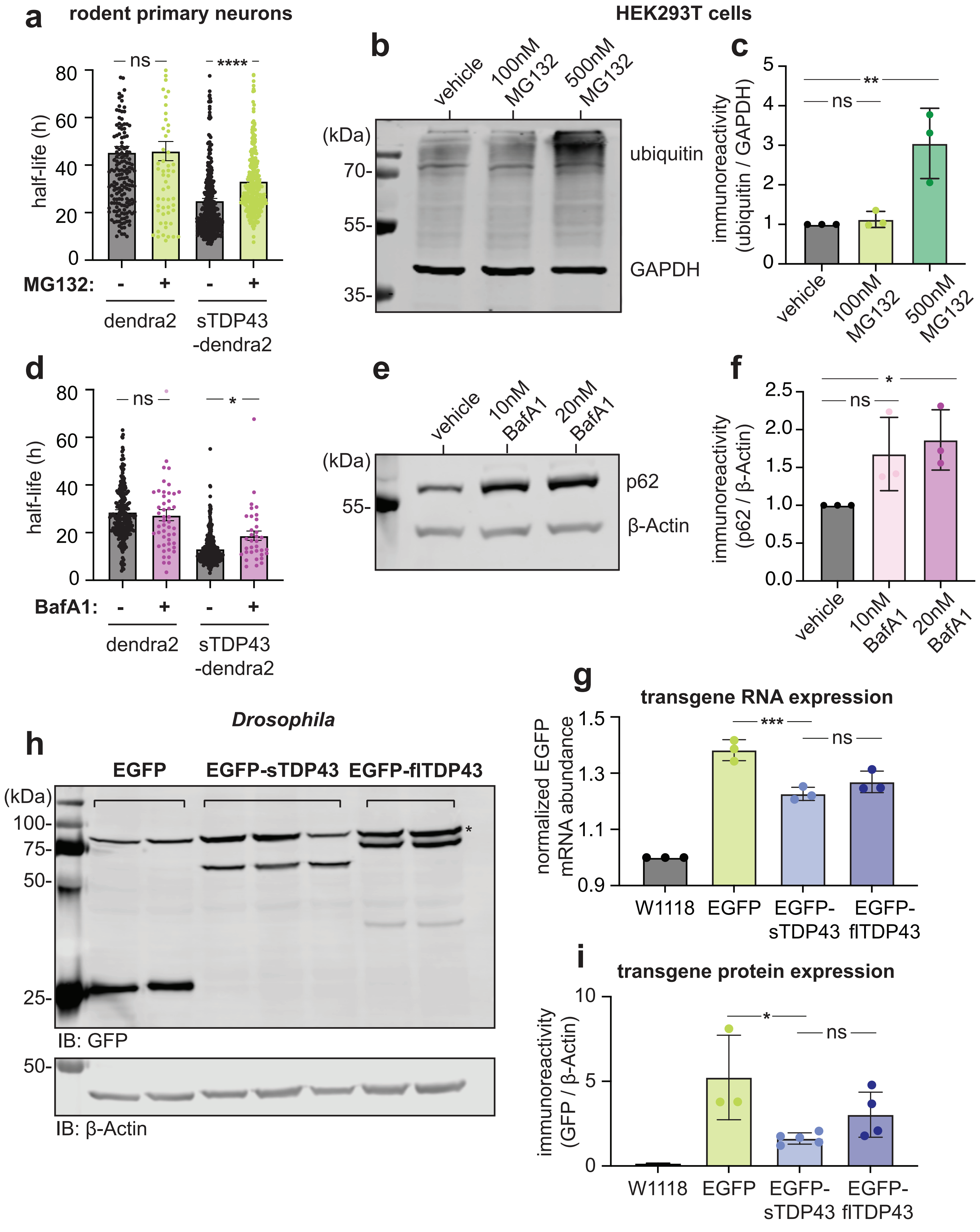

### Supplemental Figure 4

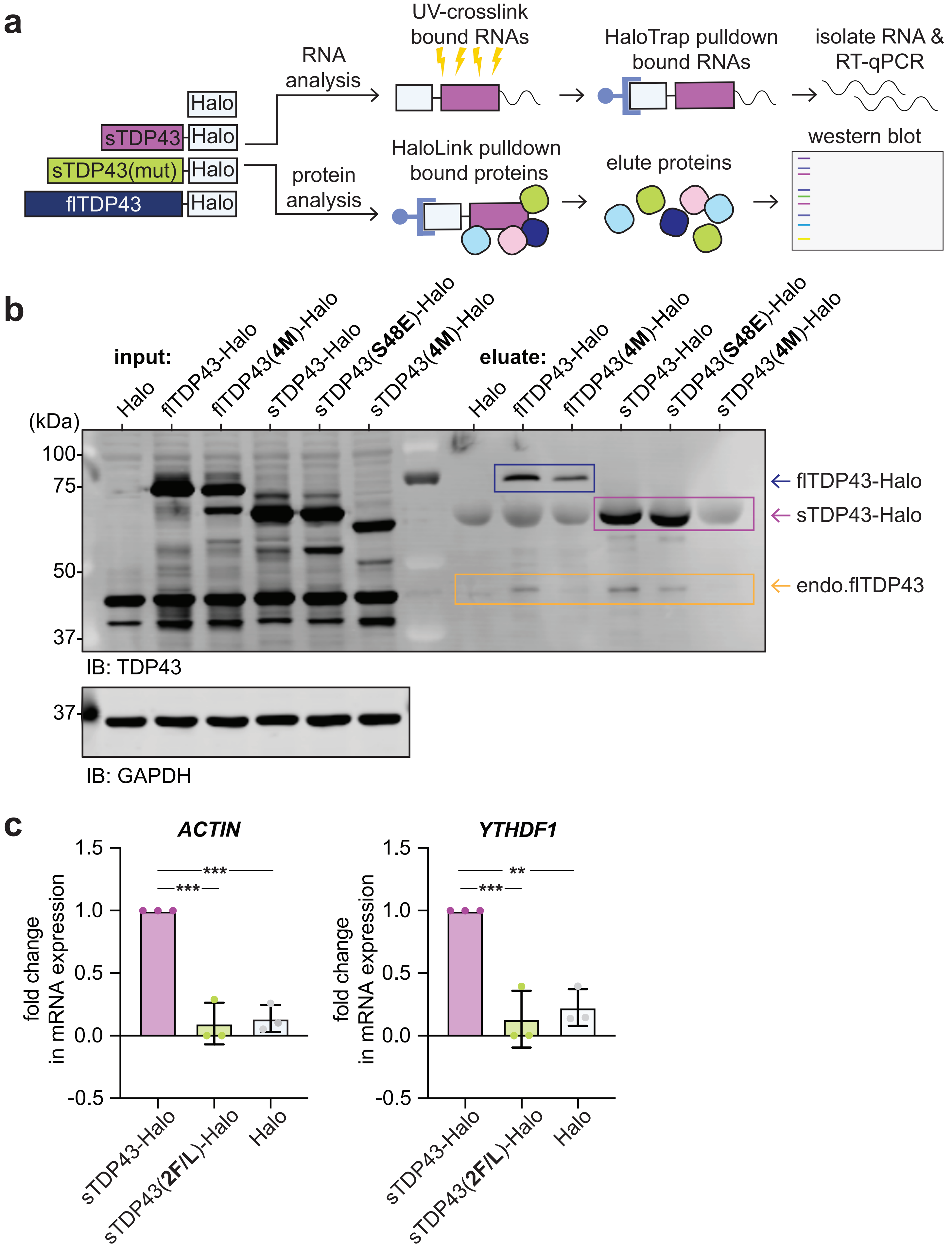

### Supplemental Figure 5

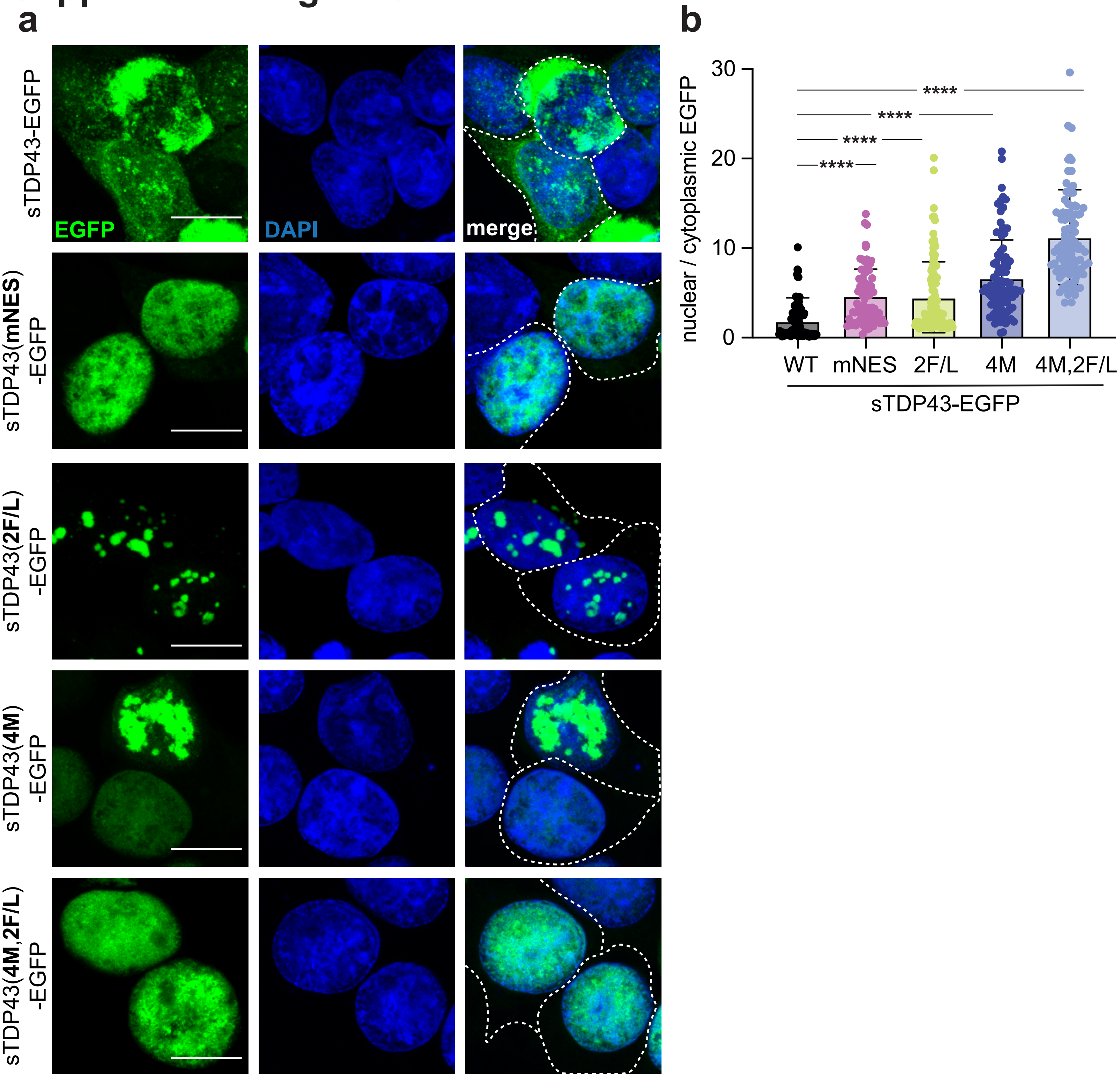

### Supplemental Figure 6

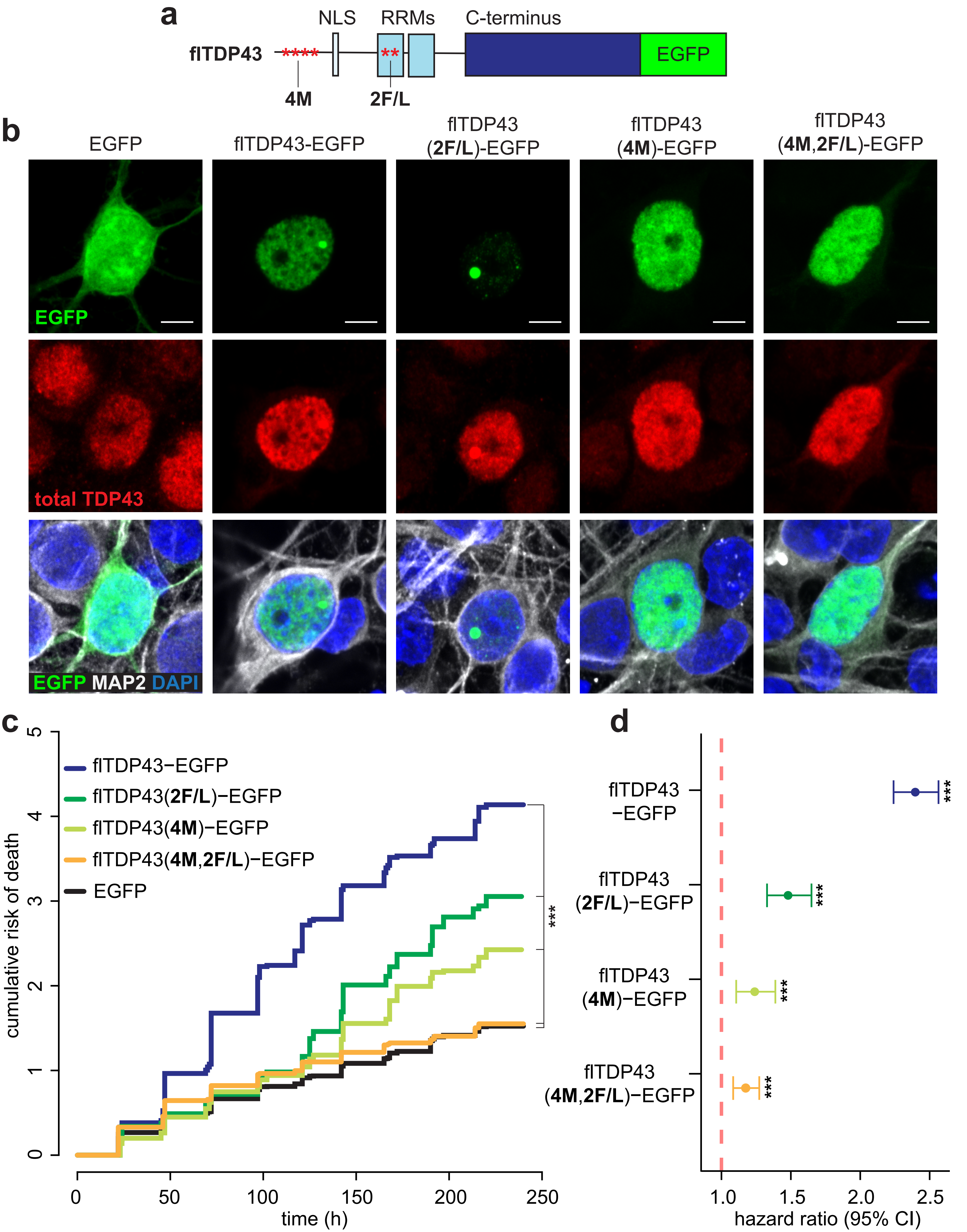

### Supplemental Figure 7

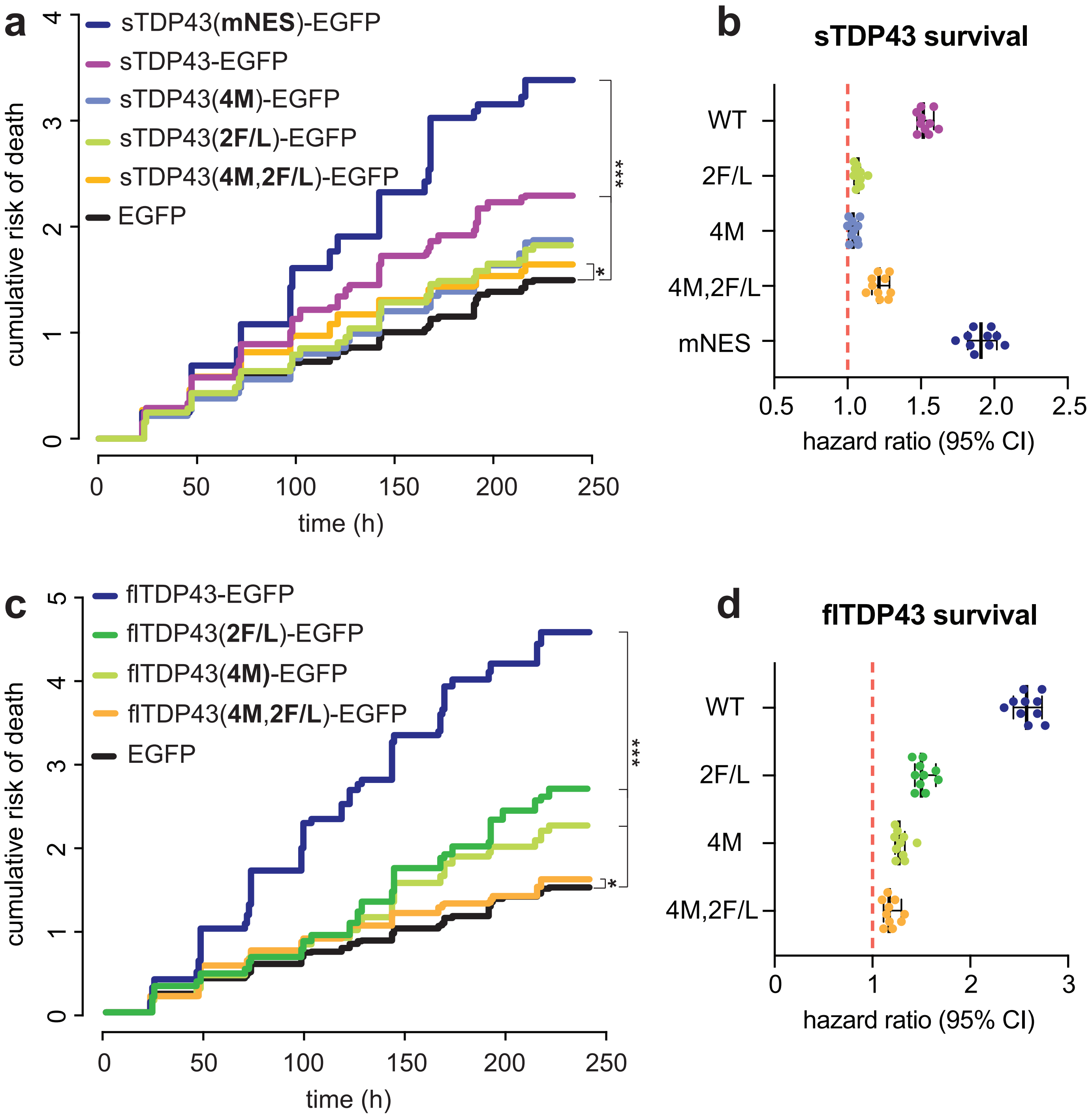

### Supplemental Figure 8

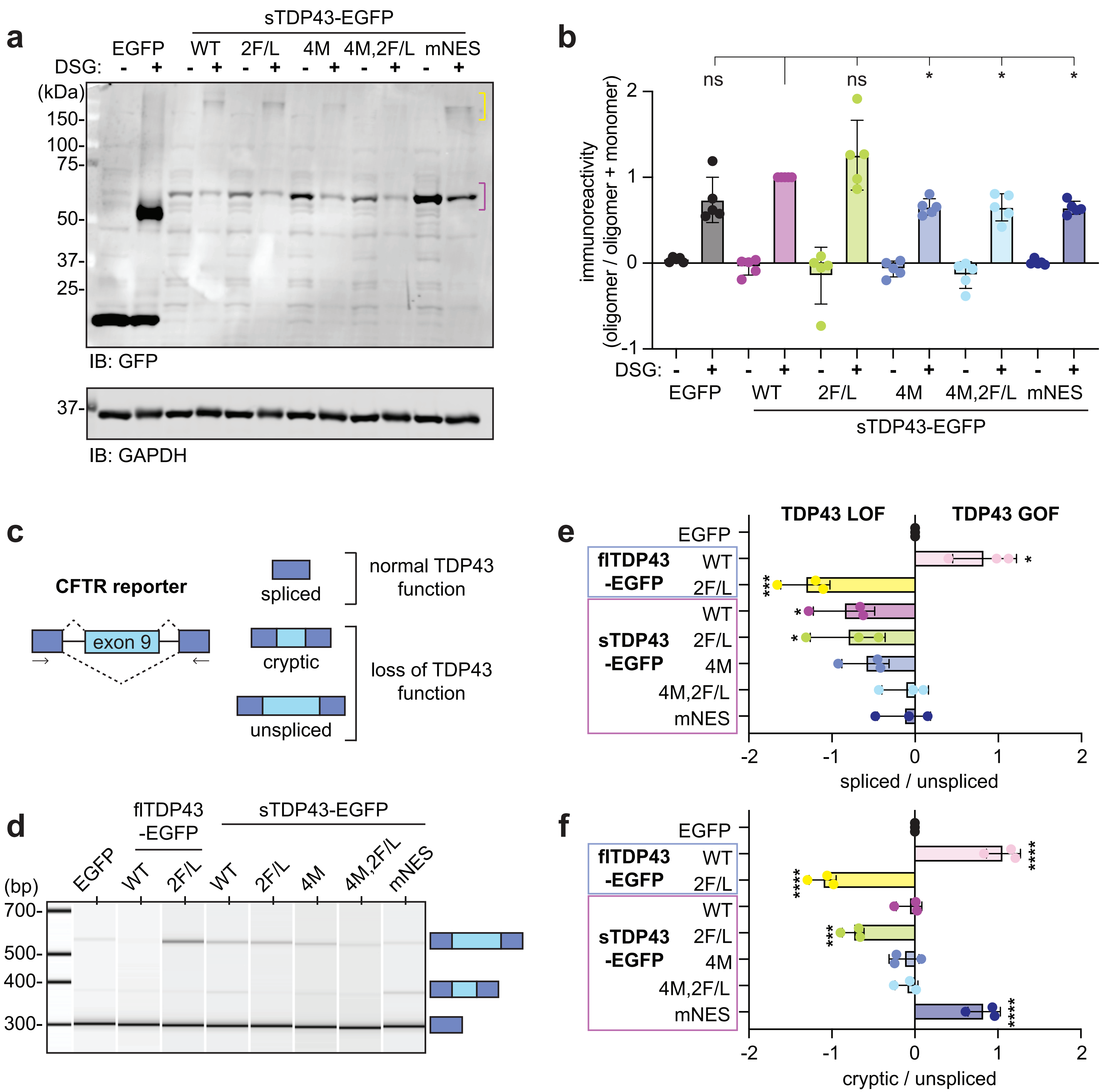
